# Cooperative DNA base and shape recognition by the CCAAT-binding complex and its bZIP transcription factor HapX

**DOI:** 10.1101/2021.07.15.452465

**Authors:** Eva M. Huber, Peter Hortschansky, Mareike T. Scheven, Matthias Misslinger, Hubertus Haas, Axel A. Brakhage, Michael Groll

## Abstract

The heterotrimeric CCAAT-binding complex (CBC) is a master regulator of transcription. It specifically recognizes the CCAAT-box, a fundamental eukaryotic promoter element. Certain fungi, like *Aspergilli*, encode a fourth CBC-subunit, HapX, to fine-tune expression of genes involved in iron metabolism. Although being a basic region leucine zipper with its own DNA recognition motif, HapX function strictly relies on the CBC. We here report two crystal structures of the CBC-HapX complex bound to DNA duplexes with distinct sequence and position of HapX sites. In either structure, a HapX dimer targets the nucleic acid downstream of the CCAAT-box and the leash-like N-terminus of the distal HapX subunit interacts with CBC and DNA. *In vitro* and *in vivo* analyses of HapX mutants support the structures, highlight the complex as an exceptional major and minor groove DNA binder, and enrich our understanding of the functional as well as structural plasticity of related complexes across species.

## INTRODUCTION

The heterotrimeric CCAAT-binding complex (CBC) is a histone-like transcription factor that is conserved from yeast to man (Brunk & Martin, 2019; Mantovani, 1999). It recognizes the CCAAT-box in the promoter region of target genes and depending on interaction partners either activates or inhibits their expression (Ceribelli et al., 2008; Hortschansky et al., 2017). By controlling a plethora of eukaryotic genes (Gurtner et al., 2017; Maity, 2017), the complex is of crucial importance (Bhattacharya et al., 2003; Oldfield et al., 2019; Yoshioka et al., 2007). Its two histone-like subunits HapC and HapE bend the target DNA in a nucleosome-like manner and provide a docking platform for HapB that recognizes the CCAAT motif (Huber et al., 2012; Nardini et al., 2013).

In certain fungi, for example *Aspergillus sp*., a subset of CBC target genes, which are involved in adaptation to iron deficiency as well as iron excess, requires the association of a fourth factor, HapX (Gsaller et al., 2014; Hortschansky et al., 2007). Based on mutagenesis and structure predictions, this subunit can be subdivided into several conserved functional segments. The C-terminal part of HapX encodes four cysteine-rich regions that are predicted to bind iron-sulfur clusters and sense intracellular iron concentrations (Hortschansky et al., 2017; Misslinger et al., 2019). Furthermore, HapX encodes a basic region (BR) leucine zipper (bZIP)/coiled-coil domain known of stand-alone transcription factors of the activator protein 1 (AP-1) family (Chen et al., 1998; Glover & Harrison, 1995). Similar to AP-1, HapX forms a homodimer and features its own DNA recognition motif (Gsaller et al., 2014). The HapX recognition signature is located about 12 base pairs (bp) downstream of the CCAAT-box, is highly promiscuous in sequence and shares only residual similarities to palindromic sequences that are usually gripped by bZIP transcription factors (T. Furukawa et al., 2020; Miller, 2009). Consistently, HapX poorly binds DNA on its own and essentially requires the core CBC as an interaction partner for high affinity promoter recognition (T. Furukawa et al., 2020). The stoichiometry of the CBC-HapX-DNA complex was determined to 1:2:1 (Hortschansky et al., 2015) and the structural motifs mediating the interaction of HapX with the core CBC were mapped to the N-terminal Hap4-like (Hap4L) domain of HapX, sharing high sequence similarity with the Hap4 protein from *Saccharomyces cerevisiae*, as well as to the αN helix of HapE (Bourgarel et al., 1999; Hortschansky et al., 2007; McNabb & Pinto, 2005). These results however raised questions about the 3D structure of the CBC-HapX-DNA complex and how the CBC can associate with two HapX subunits although providing only one HapX binding site in subunit HapE.

We here tackled this issue using X-ray crystallography and a set of biochemical techniques. Although *Aspergillus fumigatus* (Afu) is a role model for fungal virulence and pathogenicity mechanisms, including CBC-HapX controlled iron metabolism, the closely related *Aspergillus nidulans* (An) CBC crystallized more readily in previous attempts (Hortschansky et al., 2020). We therefore determined the structures of the CBC-HapX complex from *A. nidulans* bound to 35 base pairs (bp) long DNA fragments from both *A. nidulans* and *A. fumigatus*. The results highlight HapX as a major and minor groove binder and provide explanation for its low sequence specificity as well as the variable spacing between the CCAAT and HapX recognition signature. Structure-based mutagenesis and *in vivo* studies with *A. fumigatus* underpin the general relevance of the data and clarify how HapX as well as related transcription factors from yeast, *i.e.* Hap4/Pap1/Yap1 proteins, mediate their function as modulators of gene transcription.

## RESULTS

### Crystallization of the CBC-HapX-*cccA* complex from *A. nidulans*

According to our former structural studies on the CBC (Hortschansky et al., 2020; Huber et al., 2012), the core domains of the subunits HapB, HapC and HapE were unchanged (Figure S1A). Concerning HapX, we used the N-terminal part, including the CBC-binding Hap4L domain, the basic region and the coiled-coil segment. To prevent oxidation of HapX we replaced Cys92 by Lys (as in AfuHapX, Figure S1B) and exchanged Cys129, which is dispensable for HapX function (Gsaller et al., 2014), by Ser. DNA duplexes were derived from the promoter sequence of the An*cccA* gene encoding the vacuolar iron importer CccA, the strongest target of the CBC-HapX complex identified so far (Gsaller et al., 2014). Using surface plasmon resonance (SPR) experiments, the minimum length of DNA fragments required for high-affinity CBC-HapX binding was determined to about 35 bp (Figures 1 and S2). With these conditions as a guideline, more than 20 different AnCBC-HapX-An*cccA* complexes were reconstituted. DNA fragments of 35 to 42 bp with either blunt or 5’ AA/TT sticky ends were combined with AnHapX versions of different length (residues 36-133 or 36-141). Final polishing of AnCBC-HapX-An*cccA* samples by preparative size exclusion chromatography (SEC) usually yielded a single peak corresponding to the quaternary complex in a 1:2:1 CBC-HapX-DNA stoichiometry (Figure S3A). Eventually, we solved the structure of the AnCBC-HapX^36-133^ complex bound to a 35 bp long An*cccA* promoter derived DNA duplex with 5’ AA/TT overhangs to 3.4 Å resolution (PDB ID 7AW7; Figure 2A and Table S1). This DNA fragment encodes the CCAAT-box and a consensus HapX binding site (here TTAC sequence) on the Watson strand and both motifs are separated by 13 bp (Figure 3A).

**Figure 1.**
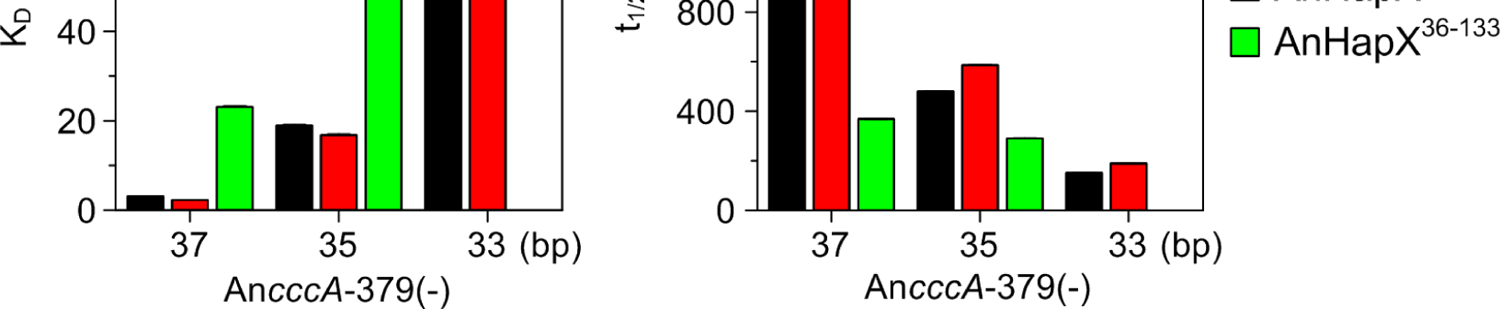
Determination of the minimal DNA duplex length and AnHapX protein size for cooperative CBC-HapX binding. SPR co-injection analysis of AnHapX binding to preformed AnCBC-DNA complexes. Dissociation constants (A) and half-lifes (B) of ternary AnCBC-DNA-HapX complexes are plotted against the DNA duplex length and were obtained by concentration dependent co-injection of HapX on preformed binary CBC-DNA complexes (two measurements per concentration). Standard deviations are indicated. Source data including errors as numbers are shown in Figure S2.

**Figure 2.**
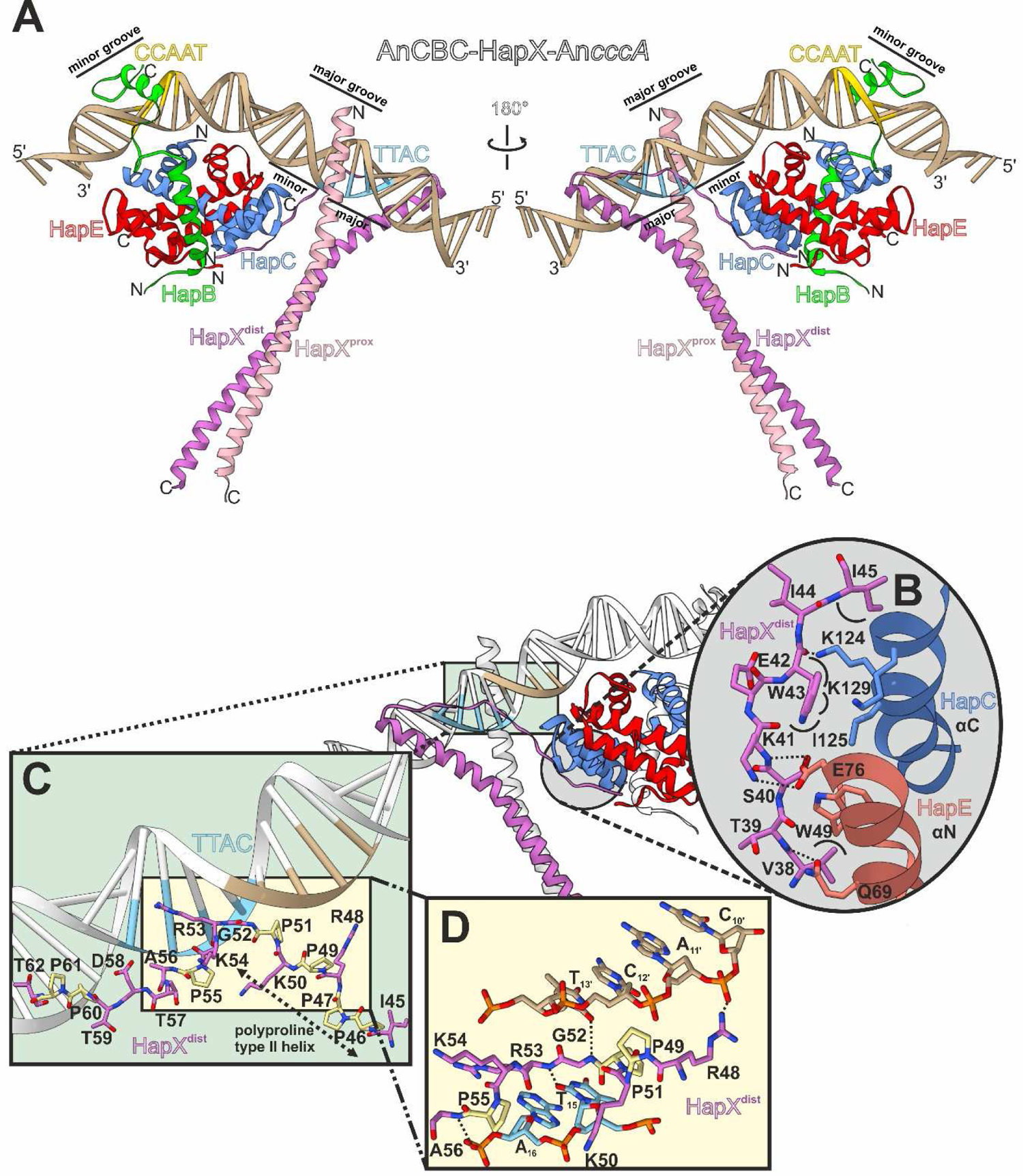
Structure of the CBC-HapX complex bound to double-stranded DNA derived from the *cccA* promoter. Protein and DNA sequences originate from *Aspergillus nidulans* (An). (A) Ribbon illustration of the multi-subunit complex. Protein chains as well as their N- and C-termini are labelled. DNA recognition motifs of HapB (CCAAT, yellow) and HapX (TTAC, lightblue) in minor and major grooves are depicted. The two HapX subunits are denoted as proximal (prox) and distal (dist) according to their relative location to the CBC on DNA. (B) The very N-terminal part of HapX^dist^ contacts the core CBC subunits HapC and HapE. Hydrogen-bonds are depicted as black dotted lines and hydrophobic contacts as black hemicycles. (C) Interactions of the HapX^dist^ N-terminus with the DNA minor groove involve a proline-rich segment (Pro residues colored yellow) of which residues 46-51 adopt a polyproline type II helix. (D) Close-up view of residues 48-56 and surrounding DNA bases. See also Figure S7A-B, G.

**Figure 3.**
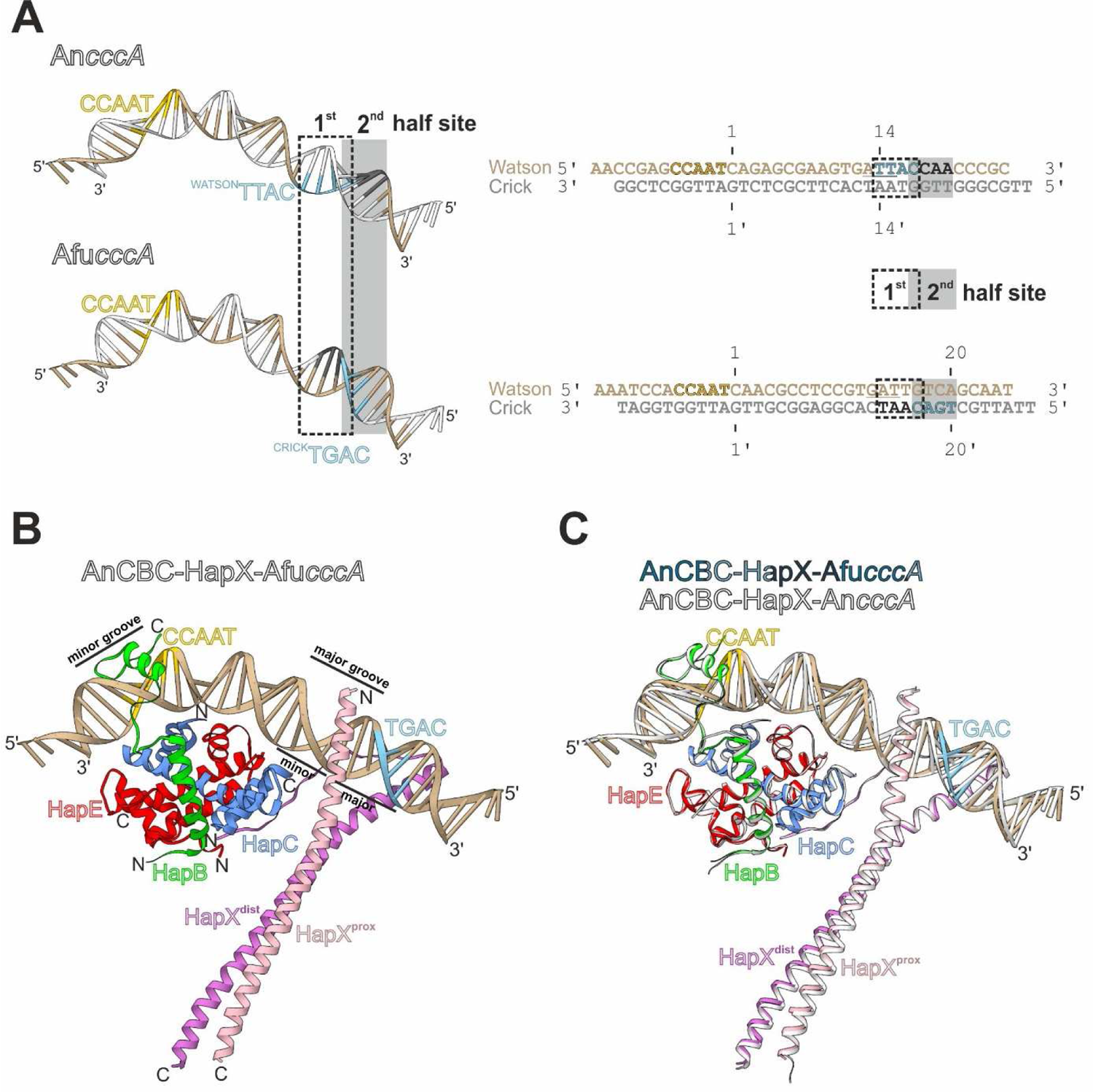
Location of the HapX binding site relative to the CCAAT-box. (A) *cccA* promoter-derived DNA fragments crystallized. For An*cccA* both CCAAT (yellow) and TTAC (TTAn; lightblue) motifs are found on the Watson strand (brown), while for Afu*cccA*, the HapX recognition site TGAC (TGAn; lightblue) is located on the Crick strand (light gray). Although the consensus HapX sequence motifs TTAn/TGAn differ in their distance to the CCAAT-box, the positions of the 1^st^ (-- boxed) and 2^nd^ (gray box) half-site of the asymmetric pseudo-palindromic recognition motif as well as the site of HapX insertion are fixed. Underlined nucleotides represent the AT-rich region (RWT submotif). (B) Ribbon illustration of the *A. nidulans* (An) CBC-HapX complex bound to double-stranded DNA derived from the *A. fumigatus* (Afu) *cccA* promoter according to Figure 2A. (C) Superposition of this chimeric complex (AnCBC-HapX-Afu*cccA*) with the AnCBC-HapX-An*cccA* structure illustrates their high structural similarity.

### Overall complex structure

In agreement with biochemical data (Hortschansky et al., 2015), one DNA double strand is occupied by one CBC core complex and two HapX subunits. Binding of the core CBC to the CCAAT-box is identical in the absence and presence of HapX (Figure S4A; r.m.s.d. 0.505 Å over 219 C_α_ atoms). The DNA is strongly bent and HapB undergoes sequence-specific interactions with nucleotides (nt) of the CCAAT motif in the minor groove (Huber et al., 2012). About 13 bp downstream, the HapX subunits are bound as a dimer to the major groove. The subunit located more distantly from the CBC on the DNA is termed HapX^dist^, whereas the one proximal to the core complex is named HapX^prox^ (Figure 2A). The HapX subunits as well as the CBC feature positively charged surfaces at their DNA binding sites that promote the association with the nucleic acid *via* electrostatic interactions (Figure S4B).

### The three-domain structure of HapX

Each HapX monomer can be subdivided into three structural segments: a C-terminal coiled-coil dimerization unit and a BR DNA binding site, which together form the bZIP motif, as well as an N-terminal Hap4L domain.

Similar to bZIP transcription factors of the AP-1 family (Chen et al., 1998; Glover & Harrison, 1995), the two HapX subunits form a coiled-coil of about 70 amino acids per chain and induce only little DNA bending of 5.8° (82.3° for AnCBC-HapX-An*cccA* versus 76.5° for AnCBC-An*cccA* (PDB ID 6Y37 (Hortschansky et al., 2020)). The helical HapX segments superimpose well (rmsd 1.457 Å over 60 C_α_ atoms), but show some structural deviation at the termini (Figure S5A). The interactions of the two HapX monomers are mostly of hydrophobic character (Figure S5B) and the mutant residues Lys92 and Ser129 are solvent-exposed (Figure S5C). Surprisingly, C-terminal extension of the crystallized coiled-coil segment by residues 134-141 enhances DNA binding up to 3.6 fold (Figure S2). According to the structural data, the increase in affinity results rather from further stabilization of the HapX homodimer than direct DNA interaction.

The BR domain of each HapX monomer inserts into the DNA major groove by interacting with both bases and the sugar-phosphate backbone (Figures 2A, S6, S7 and S8A). Notably, at a resolution of 3.4 Å, some amino acid side chains and their contact sites show defined electron density, while others are disordered and can only be supposed to mediate interactions according to their main chain tracing (Figure S7). For example, Gln72 of both HapX subunits recognizes bases located either within the TTAC binding signature on the *cccA* promoter or the corresponding non-consensus half-site of the pseudo-palindromic recognition sequence (Figures 3A, S6, S7E-F and S8A). Hydrogen bonds between Arg69 and phosphate groups are clearly discernable for HapX^dist^ and to a certain extent also for HapX^prox^ (Figures S6, S7E-F and S8A). In addition, Arg73 of HapX^dist^ and Gln67 as well as Arg78 of HapX^prox^ are targeting the sugar-phosphate backbone. Side chains Lys63, Arg64, Lys65, Arg76, Arg78 and Arg79 of HapX^dist^ and Arg64, Lys65 as well as Arg76 of HapX^prox^ are disordered but might also contribute to DNA binding – mostly *via* non-specific backbone interactions. Strikingly, the N-terminal region (residues 36-60, including the Hap4L domain) is disordered for HapX^prox^, whereas it is defined in HapX^dist^ except for two N-terminal residues (Figure 2A). Its loosely folded loop-like structure mediates crucial contacts with the core CBC as well as the nucleic acid and can be subdivided into three functional segments (Figures 2B-D and 6): The CBC binding unit (residues 39-45) of HapX^dist^ tightly interacts with the C-terminal αC helix of HapC and the N-terminal αN helix of HapE both *via* polar and hydrophobic contacts. In this regard, Trp43 of HapX^dist^ has a key function in stacking up against HapC and HapE (Figure 2B) and its well-defined electron density map served for the sequence assignment in the N-terminal region (Figure S7A). Amino acids 46-51 of HapX^dist^ form a polyproline type II helix and together with the successional residues interact with the DNA minor groove upstream of the TTAC recognition site (Figures 2C-D and S7B). Besides clear non-specific interactions of Arg48 and Ala56 with the sugar-phosphate backbone, sequence-specific contacts of the main chain atoms Gly52NH and Arg53NH of HapX^dist^ with DNA bases T_13’_ and T_15_ are assumed according to their overall position in the electron density. Moreover, the side chain of Arg53 might interact with A_16_ within the TTAC motif (Figures 2C, S7B and S8A).

In conclusion, the CBC:HapX complex represents an exceptional transcription factor that targets DNA minor and major grooves at three main sites: 1) Subunit HapB inserts into the minor groove at the CCAAT-box (Huber et al., 2012); 2) The BR domain of HapX targets the DNA major groove about 13 bp downstream of the CCAAT-box and 3) the HapX^dist^ N-terminus contacts the minor groove in between (Figure 2A). Due to these multiple interactions, association of HapX with the CBC essentially requires the presence of target DNA.

### Promiscuity of the HapX recognition motif and its structural implications

Because the HapX recognition signature is variable in sequence and position relative to the CCAAT-box (T. Furukawa et al., 2020), we aimed for additional structural data with the HapX binding site located on the Crick instead of the Watson strand (Figure 3A). To this end, we crystallized the AnCBC-HapX complex with a 35 bp long DNA double strand derived from the *cccA* promoter of *A. fumigatus* and determined its structure at 3.5 Å resolution (PDB ID 7AW9; Figures 3A-B, S3B and Table S1). This DNA fragment features a CCAAT-box on the Watson strand and a TGAC HapX consensus binding site on the opposite Crick strand at position 20. Despite this distinct structural arrangement of CBC and HapX binding motifs, we found that the CBC-HapX complex adopts the same overall structure as with the An*cccA*-derived DNA (r.m.s.d. 0.425 Å over 339 C_α_ atoms; Figure 3C). Notably, even most protein-DNA contacts are unchanged or only slightly altered (Figure S8). As with An*cccA*, only few sequence-specific DNA base interactions are observed. This is not a matter of low resolution but rather an inherent feature of HapX that also explains the low sequence conservation of the HapX binding site. The similar mode of binding to both DNA fragments is reasonable, when considering that HapX does not only recognize a four base stretch but a DNA double strand of seven nt that is, though barely related to palindromic motifs of common bZIP transcription factors, conserved in position (Figure 3A and Figure S8).

### In vitro studies on mutant A. fumigatus HapX

To validate the structural data and to underpin their universality for *Aspergilli sp.*, we conducted mutagenesis experiments with *A. fumigatus* HapX, which is crucial for virulence and antifungal drug resistance (Gsaller et al., 2016; Hortschansky et al., 2020; Schrettl et al., 2010). We selected the N-terminal Hap4L domain for mutagenesis because of its exceptional structure and so far poorly characterized dual function in CBC and DNA interaction. In particular, we deleted the CBC-interacting residues 36-42 of AfuHapX (ΔT36-I42; corresponding to amino acids 39-45 of AnHapX) to examine their functional relevance. In addition, we chose the AfuHapX residues Lys38, Trp40, Arg45 and Arg50, which are positionally equivalent to AnHapX Lys41, Trp43, Arg48 and Arg53, respectively, for point mutagenesis. Based on the X-ray structures, the lysine and arginine residues are supposed to be involved in CBC or DNA binding, respectively and the tryptophan is engaged in numerous hydrophobic contacts with the core CBC (Figure 2B-D). Using SPR, we determined the binding affinities of the mutant *A. fumigatus* HapX^24-158^ variants ΔT36-I42, K38E, W40A, R45A and R50A to three distinct preformed AfuCBC-DNA complexes. This longer HapX fragment is the reference protein that served for the determination of all our previously published HapX SPR data. Consistent with the complex structure, removal of the CBC binding portion of the Hap4L domain (ΔT36-I42) resulted in a strong decrease of cooperative HapX binding affinity to *cccA* and *cyp51A* promoter sites and binding to the *sreA* site was completely lost. In contrast, affinity of HapX ΔT36-I42 for CBC-free DNA was not affected (Figure S9). Notably, all amino acid point mutations significantly increased the dissociation constant (K_D_) of HapX (up to 40 fold) and reduced half-lifes of ternary CBC-HapX-DNA complexes (up to 8 fold; Figures 4 and S10). However, the extent was dependent on the mutation and the target DNA studied, suggesting that binding and affinity of HapX slightly vary between different promoter regions and sequences.

**Figure 4.**
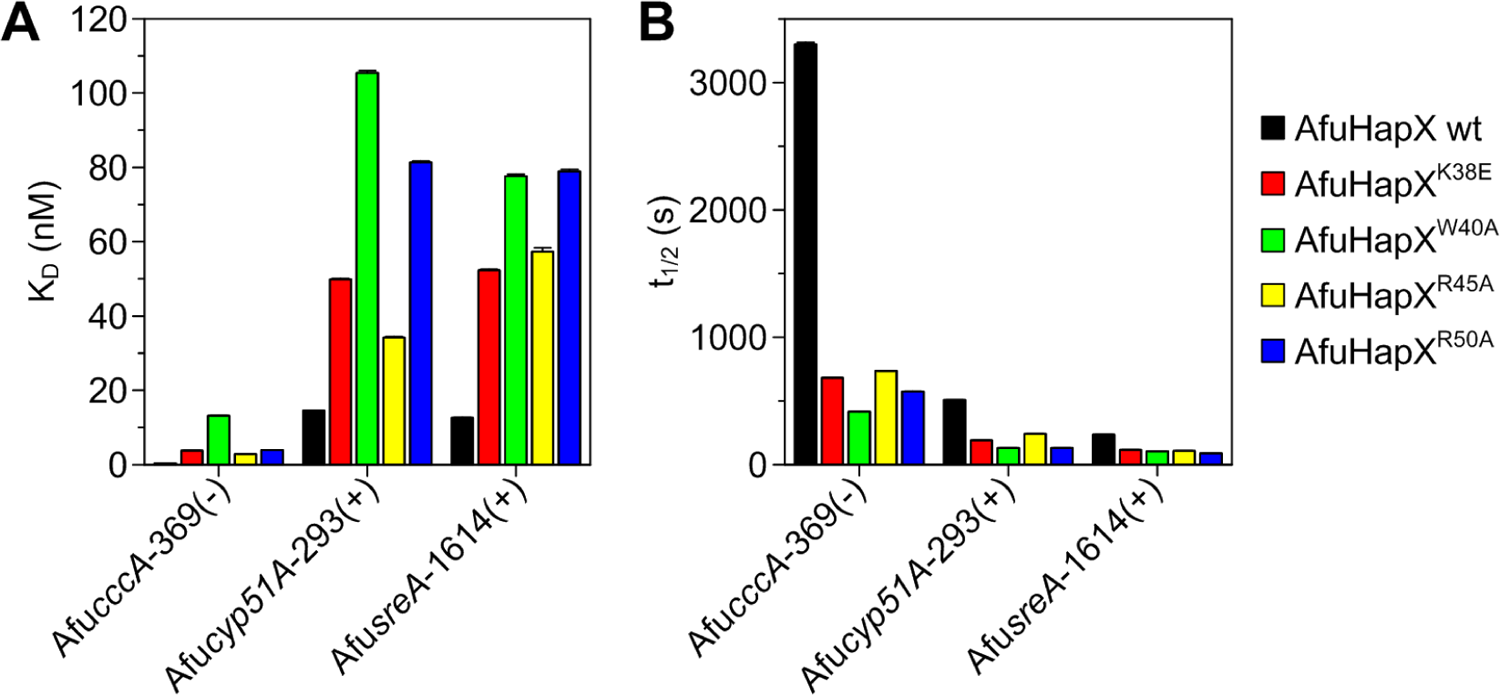
*In vitro* SPR analysis of AfuHapX Hap4L domain mutants. (A) Binding affinity of AfuHapX^24-158^ to preformed AfuCBC-DNA complexes is perturbed by mutations within the Hap4-like CBC-binding domain of HapX. (B) K38E, W40A, R45A and R50A amino acid exchanges decrease the half-lifes of the AfuCBC-DNA-HapX complex. Dissociation constants and half-lifes of ternary AfuCBC-DNA-HapX complexes are plotted over the respective DNA duplex and were obtained by concentration dependent co-injection of HapX on preformed binary CBC-DNA complexes (two measurements per concentration). Standard deviations are indicated but very small. Source data including errors as numbers are shown in Figure S10.

### In vivo mutagenesis of A. fumigatus HapX

Although being conserved and located at functionally relevant sites in the structures, the exact role of the AfuHapX Hap4L domain residues Lys38 and Arg50 remained elusive. To prove their *in vivo* functions, the mutations K38E and R50A were further evaluated in *A. fumigatus*. In addition, the mutant W40A was included, as it showed the strongest effects in the *in vitro* analysis. Potential deleterious effects of the mutations on the nuclear localization of HapX were excluded by tracking the mutant HapX proteins *in vivo* by epifluorescence microscopy (T. Furukawa et al., 2020). To this end, mutants were created in an *A. fumigatus* Δ*hapX* strain expressing a HapX version with an N-terminal Venus fluorescent protein (*^venus^hapX*).

Being a versatile regulator of adaptation to high and low iron concentrations (Gsaller et al., 2014), HapX mutants were evaluated in detail for their functionality under different iron concentrations and on several levels, *i.e.* overall fungal growth and their potential to inhibit or activate gene expression. As supposed from the *in vitro* studies and the X-ray structures, growth of *^venus^hapX^K38E^, ^venus^hapX^W40A^* and *^venus^hapX^R50A^* mutants on solid medium was impaired under iron deprivation and overload compared to wild type (wt) or *^venus^hapX* (Figure 5A). Likewise, iron starvation and excess conditions caused growth retardation of the mutant strains in liquid cultures (Figure 5B). In agreement, under low iron concentrations and compared to control strains, K38E, W40A and R50A mutants were less effective in repressing iron-consuming pathways such as production of heme proteins (*cycA*) and heme biosynthesis (*hemA*) (Figure 5C). Despite this functional impairment, the mutants did not behave like Δ*hapX* (Figure 5C) or a strain that lacks the entire Hap4L domain (T. Furukawa et al., 2020). Furthermore, we found that HapX-mediated activation of genes required for the biosynthesis and uptake of endogenously produced iron chelators (siderophores) like triacetylfusarinine C (TAFC) was drastically reduced in all mutants. In particular, mRNA levels of the HapX target genes *sidG* and *mirB*, encoding an acetyltransferase involved in TAFC biosynthesis and a siderophore transporter for re-uptake of iron-loaded TAFC, respectively (Schrettl et al., 2010), were markedly lower (Figure 5D). In addition, total production of TAFC was limited to about 20% of the wt level (Figure 5E). These results indicate that the mutations studied affect both activation and inhibition of gene expression by HapX.

**Figure 5.**
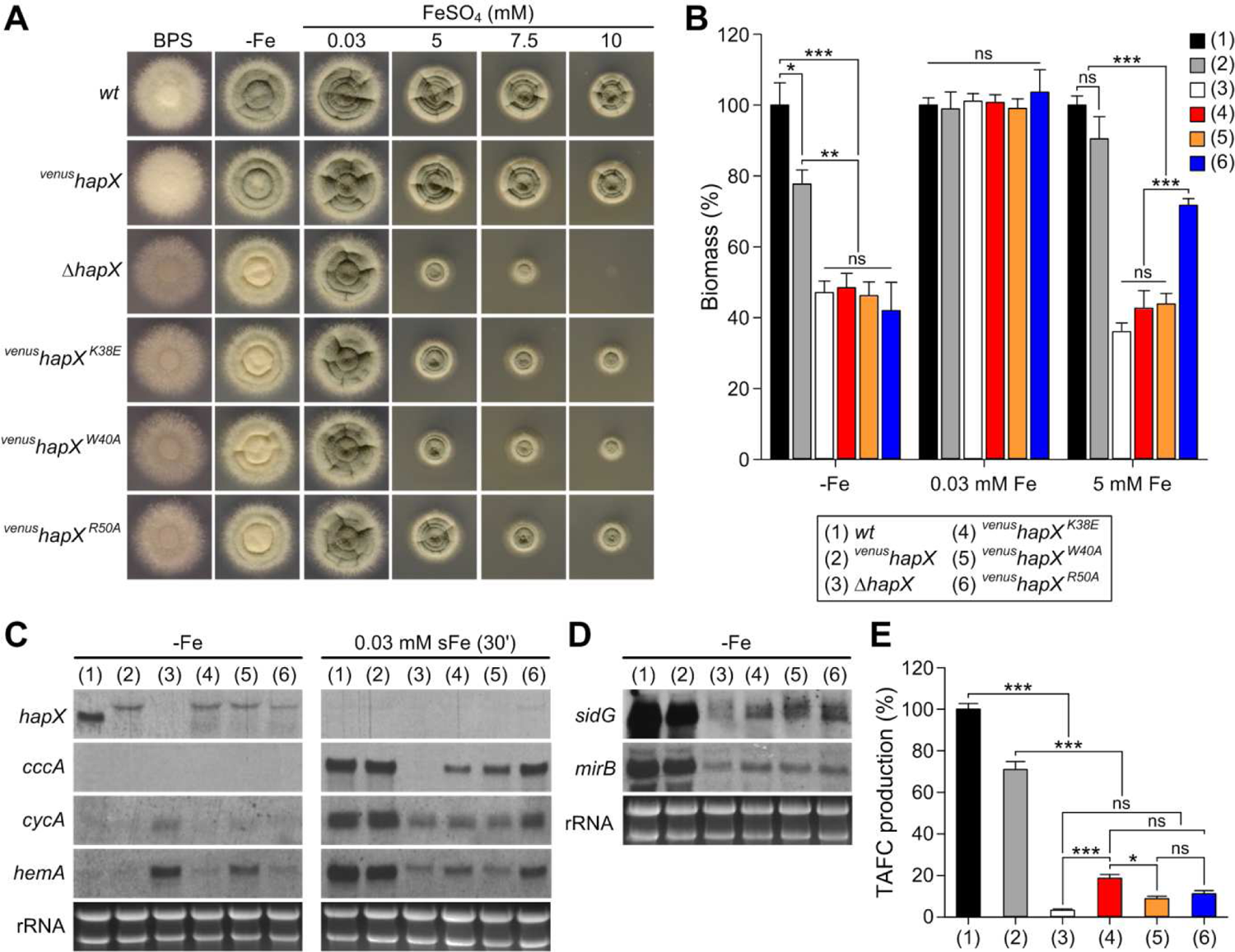
K38E, W40A and R50A mutations impair AfuHapX functions *in vivo* during both iron starvation and iron excess. (A) Growth pattern of *A. fumigatus* wild type (*wt*), *^venus^hapX*, Δ*hapX* and mutant strains expressing *^venus^hapX* alleles carrying either K38E, W40A or R50A amino acid substitutions on solid minimal medium at 37 °C for 48 h. (B) Production of biomass in submersed cultures during iron starvation (-Fe), iron sufficiency (0.03 mM FeSO_4_, +Fe) and iron excess (5 mM FeSO_4_, hFe) after liquid growth at 37 °C for 24 h. Biomass production of the mutant strains was significantly lower compared to *wt* and *^venus^hapX* during -Fe and hFe, but not +Fe. (C) CBC-HapX interaction (Lys38, Trp40) and HapX DNA-binding (Arg50) are essential for the gene regulatory function during iron starvation and short-term iron excess. Northern blot analyses were performed after liquid growth for 18 h under iron starvation (-Fe) and from mycelia shifted for another 30 min from -Fe to iron sufficiency (0.03 mM FeSO_4_, sFe). rRNA levels are shown below as loading control. (D) Lys38, Trp40 and Arg50 of HapX are crucial for activation of genes that promote TAFC siderophore biosynthesis and uptake in *A. fumigatus*. (E) Production of TAFC in the absence of iron is impaired by the amino acid residue substitutions K38E, W40A and R50A. Data are presented as the mean and standard deviation of three biological replicates, and analyzed by one-way ANOVA with Tukey’s multiple comparison test (*, *P* ≤ 0.05; **, *P* ≤ 0.01; ***, *P* ≤ 0.001; ns, not statistically significant). Source data are provided as separate files.

**Figure 6.**
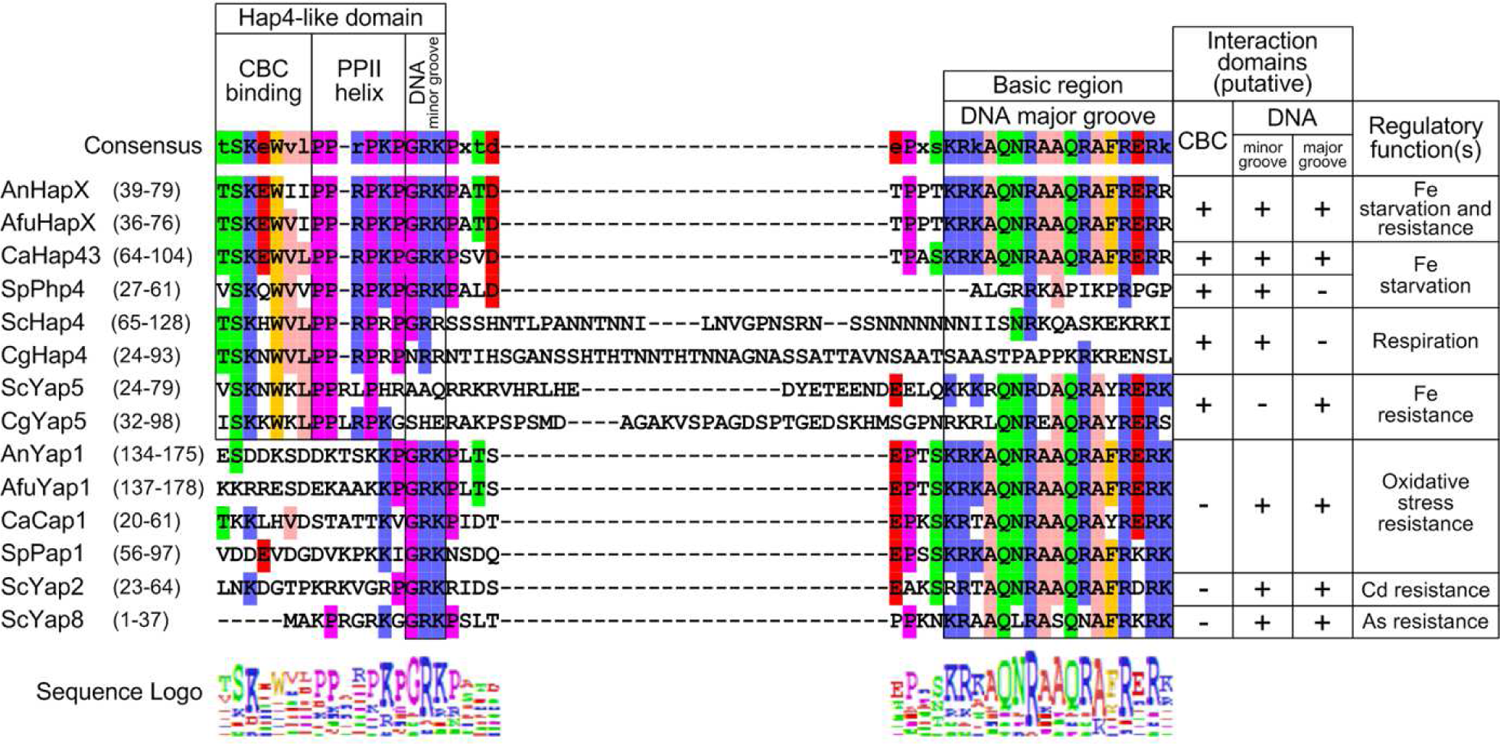
Modules of the multifunctional Hap4L domain are present in iron regulators of the HapX-type and in AP-1 bZIP proteins of the Yap stress response family. Alignment of the Hap4L domain and basic region of HapX as well as AP-1 proteins from *A. nidulans* (An), *A. fumigatus* (Afu), *Candida albicans* (Ca), *Candida glabrata* (Cg), *Schizosaccharomyces pombe* (Sp) and *S. cerevisiae* (Sc). Residues that match the consensus sequence are colored according to their physico-chemical properties. Alignments were generated using MegAlign Pro 17 (DNASTAR, Inc.). AnHapX, AfuHapX and its homolog Hap43 in *C. albicans* (Singh et al., 2011), are bZIPs that share the canonical 16 aa Hap4L domain. While HapX regulates adaptation to iron starvation and excess, Hap43 lacks a role in iron resistance (Skrahina et al., 2017). Php4 in *S. pombe*, which represses iron-dependent pathways in response to iron starvation, contains the Hap4L domain, but lacks a bZIP domain (Mercier et al., 2008). The same applies to the Hap4 activators of respiratory gene expression in *S. cerevisiae* and *C. glabrata* (Bourgarel et al., 1999; Thiebaut et al., 2017). In contrast, Yap5 bZIP-type activators of the high iron stress response in *S. cerevisiae* and *C. glabrata* (Merhej et al., 2015) contain a degenerated Hap4L domain lacking the DNA minor groove binding part. Notably however, this module is present in several Yap1/2 and Yap8 family members and N-terminally flanks the basic regions of these bZIPs.

To evaluate adaptation to environmental changes in iron concentration, strains were shifted for 30 min from iron starvation to sufficiency (Figure 5C). While control strains efficiently activated genes required for iron consumption under these conditions, the mutants showed significantly less induction of *cycA* and *hemA* genes. In fact, *cycA* and *hemA* expression levels were comparable to Δ*hapX*. Regarding *cccA*, induction was less affected in the K38E, W40A and R50A mutants compared to Δ*hapX* – an effect that might result from the three CBC:HapX binding sites in the *cccA* promoter (T. Furukawa et al., 2020).

In summary, under iron deprivation or short-term variations of iron concentrations, transcriptional control by the AfuHapX mutants K38E, W40A and R50A was significantly impaired. Although the effects strongly depended on the target gene and promoter-specific characteristics, all mutants showed the same trend. The fact that the point mutants did not phenocopy Δ*hapX* agrees with the SPR-data and the X-ray structures visualizing multiple protein-protein and protein-DNA contacts that can compensate single point mutations and allow HapX to find its destination on the nucleic acid yet at lower efficacy.

## DISCUSSION

### Homodimerization of HapX

bZIP transcription factors form long α helices that wrap around each other to build a superhelical coiled-coil structure. At the DNA binding site, the helices separate and bind like chopsticks to the major groove of the nucleic acid. As homo- or heterodimers, bZIP proteins bind short palindromic or pseudo-palindromic DNA sequences (Pathak & Sigler, 1992). HapX is the fourth subunit of the CBC complex in filamentous fungi and forms a homodimer. Sequence analyses of HapX subunits reveal that the leucine zipper/coiled-coil domain responsible for homodimerization of HapX is exceptional. Members of the bZIP family contain heptad repeats of leucine (or hydrophobic) residues (leucine zipper) that form an amphiphilic α-helix, whose hydrophobic side is buried upon dimerization. For HapX however an insertion of four residues is observed in heptad 3 (Figure S1B) and this peculiar feature is conserved in all known HapX subunits (Figure S11). Although experimental evidence is missing, it is tempting to speculate that this insertion may prevent heterodimerization of HapX with the numerous other bZIP factors present in fungal cells (T. Furukawa et al., 2020) and ensure productive assembly of CBC-HapX-DNA complexes.

### DNA recognition by HapX

In contrast to stand-alone bZIP transcription factors, DNA recognition elements cooperatively bound by bZIPs and additional regulators often do not exhibit optimal consensus motifs (Chen et al., 1998; Panne et al., 2004). According to its conserved amino acid signature NxxAQxxFR, HapX belongs to the Pap1/Yap1 subfamily of AP-1 proteins. Members of this family specifically recognize the two adjacent TTAn half-sites of the consensus binding sites TTAcgTAA or TTAcTAA *via* their helical bZIP domains (Fujii et al., 2000). HapX however recognizes a rather promiscuous DNA motif with usually only a single consensus half-site (Figure 3A). This half-site can be present on the Watson or Crick DNA strand and either matches the TTAn or TGAn (TKAn) recognition sequence characteristic of the Pap1/Yap1 and the Fos/Jun/Gcn4 subfamily of bZIP factors, respectively (T. Furukawa et al., 2020; Gordan et al., 2011; Gsaller et al., 2014). In addition, all CBC-HapX target sites share a clustering of adenine and thymine bases with the consensus sequence RWT (R=A/G; W=A/T) exactly 12 bp downstream of the respective CCAAT motif (T. Furukawa et al., 2020). Importantly, these RWT submotifs, here ATT (An*cccA*) and GAT (Afu*cccA*), are indispensable for the proper regulation of HapX target genes (T. Furukawa et al., 2020) and create a minor groove that is predicted to be more narrow (< 5Å) compared to ideal B-DNA (5.8 Å) (Figure S12). Narrow minor grooves are preferentially targeted by arginine residues (Rohs et al., 2009) and Arg53 of AnHapX^dist^ (Arg50 of AfuHapX^dist^) is ideally placed to do so as well. Although the side chain of Arg53 is not resolved in the electron density map (Figure S7B), its relevance for HapX function has been proven by *in vitro* and *in vivo* mutagenesis (Figures 4 and 5). Considering that the TKAn HapX recognition motif is poorly conserved in sequence and that most interactions of HapX with the nucleic acid involve the sugar-phosphate backbone (Figure S8), HapX rather recognizes local sequence-dependent DNA shape than specific nt sequences. In support of this model, shape-motifs are frequent in regions targeted by multiple transcription factors (Samee et al., 2019; Slattery et al., 2014) such as the CBC-HapX binding area that often overlaps with other protein binding sites (Gsaller et al., 2016). In conclusion, specificity and strength of cooperative CBC-HapX-DNA-binding cannot be explained *via* primary nt sequence signatures alone and involve also shape-based DNA recognition.

### Spacing and relative positions of CBC and HapX binding sites

The downstream position of HapX TTAn/TGAn half-sites relative to the CCAAT-box can vary between 11-23 bp with strong preference for 14 nt (both encoded on the Watson strand) and 20 nt (HapX binding site encoded on the Crick strand) (Figure 3A) (T. Furukawa et al., 2020). Here we present crystal structures for both cases using An*cccA* (position 14; both motifs on Watson strand) and Afu*cccA* DNA (position 20; motifs on Watson and Crick strand), respectively. Notably, for both nucleic acid fragments, the overall complex structures as well as protein-DNA interactions are identical and HapX^prox^ binds about 13 nt downstream of the CCAAT-box. Because both HapX^prox^ and HapX^dist^ bind consensus TTAn/TGAn as well as nonconsensus signatures and assemble into a homodimer, the rough position of the bZIP factor on the nucleic acid is the same irrespective of the two possible positions for the half-site motif on the DNA double strand. Distances much shorter than 13 bp are prevented by steric hindrance with the core CBC, but minor difference in spacing may be tolerated. In this regard, the polyproline type II helix of HapX^dist^, a structural element that is more flexible than α-helices (Adzhubei et al., 2013), is ideally suited to function as a leash for the bZIP domains to find their destination at the DNA. This extraordinary structural flexibility could assist control of *sreA* and *leuA* genes. In the promoters of these genes, the distance between CCAAT box and the HapX TGAC half-site, both located on the Watson strand, is larger than for most other genes, namely 20 respectively 21 nt (Figure S13) (Furukawa et al., 2020). This exceptional arrangement of recognition motifs could favor interactions of the Hap4L domain of HapX^prox^ with the DNA minor groove and the CBC, resulting in a rewired overall complex architecture.

### Interactions of HapX with the CBC and implications for related transcription factors

High-affinity DNA binding by HapX essentially requires interaction with the core CBC. Interestingly, the CBC interaction domain of HapX (including residues Lys41, Trp43, Arg48 and Arg53) shows high sequence similarity to the Hap4 proteins from yeasts (Figure 6) (Bourgarel et al., 1999; McNabb & Pinto, 2005; Sybirna et al., 2005). Hap4 triggers gene expression by associating as a monomer with the heterotrimeric Hap core complex (Hap2/3/5) (Mao & Chen, 2019). In line with findings that Hap4 binds to Hap3 (Xing et al., 1993), our structural model visualizes tight interactions between HapX and HapC, the Hap3 homologue. Furthermore, two models have been proposed previously to explain why DNA binding is necessary for stable association of Hap4 to the Hap core complex (McNabb & Pinto, 2005). The first model suggested a conformational change of the Hap complex upon DNA binding, whereas the second proposed that Hap4 targets both the DNA and the heterotrimeric Hap core complex. Our structures now clearly argue for the second version and clarify a long-standing mystery of the Hap complex in yeasts. Besides, sequence alignments based on our X-ray structures imply that transcription factors of the Pap1/Yap1 subfamily of AP-1 proteins sharing the (KP)GRK motif of the Hap4L domain may target DNA minor grooves as well (Figure 6). The structure of the Pap1-DNA complex does not visualize such a contact, because the Pap1 protein used for crystallization lacked the first three amino acids of the KPGRK motif and in addition, the N-terminal four residues of the Pap1 construct were disordered in the electron density map (Fujii et al., 2000). However, target gene activation by and the stress response function of the Pap1 family member Yap8 from *S. cerevisiae* was recently shown to depend on the corresponding KGGRK amino acid stretch (Maciaszczyk-Dziubinska et al., 2020). Notably, minor groove binding *via* the KPGRK motif is not restricted to eukaryotic bZIPs of the HapX-type. The MogR transcriptional repressor of the bacterial pathogen *Listeria monocytogenes* for example uses the KPGRK sequence to target its AT rich DNA recognition motif *via* the minor groove (Shen et al., 2009) (Figure S14).

Altogether, the here reported structures visualize the interactions of a Hap4-like domain protein with its corresponding core complex and AT-rich DNA minor grooves. The results provide evidence for a shape-based DNA recognition mechanism by HapX factors and suggest that members of the Pap1/Yap1 bZIP transcription factor family bind their target DNA similarly.

## MATERIAL AND METHODS

### Overproduction and purification of recombinant CBC and HapX

The *A. nidulans* CBC consisting of the conserved core domains of HapB, HapC, and HapE was produced and purified as described (Gsaller et al., 2014; Huber et al., 2012). Briefly, synthetic genes coding for HapB^231-293^, HapC^42-132^ and HapE^47-164^ were sequentially cloned in the pnCS vector for expression of a polycistronic transcript (Diebold et al., 2011). The expression plasmids were transformed in *Escherichia coli* BL21(DE3). After overnight autoinduction and cell lysis, the heterotrimeric CBC was purified to homogeneity by subsequent cobalt chelate affinity and size exclusion chromatography (SEC). Size exclusion fractions containing pure CBC were pooled based on SDS-PAGE analysis, concentrated by ultrafiltration (Amicon Ultra-15 10K centrifugal filter device, Millipore) to 25-30 mg/ml, aliquoted, flash frozen in liquid nitrogen, and stored at −80 °C. A synthetic gene coding for *A. nidulans* HapX^36-133M^ carrying C92K and C129S point mutations in its coiled-coil domain was expressed as an N-terminal maltose-binding protein (MBP) fusion separated by a tobacco etch virus (TEV) protease site in *E. coli* BL21(DE3), *via* a modified pET28a vector (Novagen). Crude bacterial lysates were purified by Dextrin Sepharose affinity chromatography (GE Healthcare) after overnight autoinduction and cell lysis. The MBP-HapX fusion was cleaved with TEV protease and further purified sequentially using CellufineSulfate (Millipore) affinity chromatography, (NH_4_)_2_SO_4_ precipitation (40% w/v), and SEC on a Superdex prep grade 200 26/60 column (GE Healthcare) in 20 mM Tris/HCl, 2 M NaCl, 2 mM DTT, pH 7.5. SEC fractions were concentrated by ultrafiltration (Amicon Ultra-15 10K centrifugal filter device, Millipore) to 25-30 mg/ml, aliquoted, flash frozen, and stored at −80 °C. All other *A. nidulans* HapX bZIP polypeptides used in this study (HapX^27-141^, HapX^36-141^, HapX^36-133^ or HapX^62-141^) as well as the *A. fumigatus* HapX^24-158^ wild type and mutant bZIP proteins were produced the same way.

### Surface plasmon resonance (SPR) measurements

Real-time SPR protein-DNA interaction measurements were performed according to previously published protocols (T. Furukawa et al., 2020; Gsaller et al., 2016). Notably, for cooperative CBC-HapX binding analysis measured by SPR co-injection on *A. fumigatus* promoter motifs, an *A. fumigatus* CBC consisting of the subunits HapB^230-299^, HapC^40-137^ and HapE^47-164^ was purified as described for the *A. nidulans* CBC.

### Preparation of CBC-HapX-DNA complexes for crystallization

HPLC-purified oligonucleotides were purchased (biomers), dissolved in 10 mM Tris/HCl, 50 mM NaCl, 1 mM EDTA, pH 8.0 at a concentration of 5 mM and annealed by mixing equal volumes of each strand to yield a final DNA duplex concentration of 2.5 mM. Complexes were prepared by mixing 1:2:1.2 equivalents of AnCBC, AnHapX^36-133M^ subunits and DNA at a concentration of 0.4-0.5 mM in a buffer containing 20 mM Tris/HCl, 0.9 M NaCl, 3 mM DTT, pH 7.5. CBC-HapX-DNA preparations were dialyzed (Slide-A-Lyzer Dialysis Cassette G2, MWCO 10 kDa; Thermo Fisher) against SEC Buffer (20 mM Tris/HCl, 100 mM NaCl, 1 mM DTT, pH 7.5) and further purified using a Superdex prep grade 200 16/60 column (GE Healthcare). The presence of all three CBC subunits, HapX as well as DNA in the collected main fraction was verified by a dual stain method that allows independent visualization of the protein and nucleic acid species (Pryor et al., 2012). SEC purified protein-DNA preparations were concentrated 10-fold by ultrafiltration (Amicon Ultra-15 30K centrifugal filter device, Millipore) and subjected to crystallization.

### DNA shape analysis

Sequence-dependent DNA minor groove width and electrostatic potential were determined with DNAphi (https://rohslab.usc.edu/DNAphi/index.html (Chiu et al., 2017)).

#### Crystallization and structure determination

The CBC-HapX-*cccA* complex was crystallized by the sitting drop vapor diffusion technique at 20 °C. Crystal drops (0.4 µL) contained equal volumes of the protein-DNA complex (30 mg/ml) and reservoir solution. The 35 bp long DNA sequence was derived from the promoter encoding the vacuolar iron transporter (*cccA*) of either *A. nidulans* (An) or *A. fumigatus* (Afu) and carried 5’ AA/TT overhangs. The CBC-HapX-An*cccA* complex crystallized from 0.2 M ammonium fluoride, 20% (w/v) PEG3350 and the CBC-HapX-Afu*cccA* from 0.1 M Tris/HCl pH 8.5, 25% (w/v) PEG4000. Crystals were cryoprotected by the addition of a 1:1 (v/v) mixture of mother liquor and 60% (v/v) ethylene glycol and subsequently super-cooled in liquid nitrogen. Diffraction data were collected at the beamline X06SA, Swiss Light Source, Villigen, Switzerland at λ=1.0 Å. Reflection intensities were analyzed with the program package XDS (Kabsch, 1993). The structure of CBC-HapX-An*cccA* was determined by sequential Patterson search calculations with PHASER (McCoy et al., 2007) using the coordinates of *A. nidulans* CBC in complex with a 23 bp fragment of the *cccA* promoter (PDB ID:6Y37 (Hortschansky et al., 2020)) and two chains of the human FosB bZIP domain (PDB ID 5VPE) (Yin et al., 2017). The structural model of the AnCBC-HapX-An*cccA* complex was used to phase the AnCBC-HapX-Afu*cccA* structure. Cyclic refinement and model building steps were performed with REFMAC5 (Vagin et al., 2004) and Coot 0.9 (Emsley et al., 2010). DNA restraints for model building and refinement were taken from the previously reported AnCBC-An*cccA* (PDB ID 6Y37 (Hortschansky et al., 2020)) and AfuCBC-Afu*cccA* (PDB ID 6Y36 (Hortschansky et al., 2020)) structures, respectively. TLS refinements finally yielded good values for R_crys_ and R_free_ as well as root-mean-square deviation (r.m.s.d.) bond and angle values. The models were proven to fulfill the Ramachandran plot using PROCHECK (Laskowski et al., 1993) (Table S1). The DNA bending angle was analyzed with the Curves+ and SUMR algorithms (Lavery et al., 2009). Graphical illustrations were created with the UCSF Chimera package from the Resource for Biocomputing, Visualization, and Informatics at the University of California, San Francisco (Pettersen et al., 2004). Electrostatic surface potentials were calculated with the APBS online tool available at http://server.poissonboltzmann.org/ and the results visualized with UCSF Chimera. The structures were analyzed for their minor and major groove contacts by DNAproDB (Sagendorf et al., 2017; Sagendorf et al., 2020) and for the presence of a polyproline type II helix using PolyprOnline (Chebrek et al., 2014). Coordinates and structure factors have been deposited in the protein data bank under the accession codes: 7AW7 (AnCBC-HapX-An*cccA*) and 7AW9 (AnCBC-HapX-Afu*cccA*).

### Generation of mutant *A. fumigatus* strains

*A. fumigatus* strains used in this study are listed in Table S2. For point mutations of the AfuHapX Hap4L domain, plasmid phapX^VENUS^-hph (Gsaller et al., 2014) was used as template and modified using QuikChange II site-directed mutagenesis kit (Agilent for K38E and R50A) or Q5 site-directed mutagenesis kit (New England Biolabs; for W40A). Primers used for mutagenesis are listed in Table S3A. *A. fumigatus* strain Δ*hapX* (Gsaller et al., 2014) was transformed with 5 µg each of the mutagenized plasmid, which was linearized through *Sna*BI digestion. *A. fumigatus* mutants were selected using 0.2 mg/ml hygromycin B. Correct single-copy integration at the *hapX* locus was confirmed by Southern blot analysis.

### Growth analysis of *A. fumigatus*

For growth phenotyping, Northern blot and siderophore production analysis, fungal strains were cultivated at 37 °C in *Aspergillus* Minimal Medium (AMM) with 1% (w/v) glucose as carbon source and 20 mM glutamine as nitrogen source as previously reported (T. Furukawa et al., 2020). To increase iron depletion in solid minimal medium, 0.2 mM of the iron chelator bathophenanthroline disulfonate (BPS) was added. Growth assays were performed with either 10^4^ or 10^8^ conidia inoculated on solid medium or grown in 100 ml liquid culture, respectively.

### Northern blot analysis and measurement of siderophore production

Northern blot analysis and measurement of siderophore production were conducted as previously reported (T. Furukawa et al., 2020). DIG-labeled hybridization probes were generated by PCR. Primers for amplification of hybridization probes are listed in Table S3B.

## ACCESSION NUMBERS

Coordinates and structure factors have been deposited in the protein data bank (entry codes: 7AW7 (AnCBC-HapX-An*cccA*), 7AW9 (AnCBC-HapX-Afu*cccA*); see also Table S1).

## ACKNOWLEDGMENTS

We thank Sylke Fricke for excellent technical assistance and are grateful to the staff of the beamline X06SA at the Paul-Scherrer-Institute, Swiss Light Source, Villigen Switzerland for assistance during data collection.

## FUNDING

This work was funded by the joint D-A-CH program ‘Novel molecular mechanisms of iron sensing and homeostasis in filamentous fungi’ (Deutsche Forschungsgemeinschaft (DFG) to AAB (BR 1130/14-1), MG (GR 1861/8-1) and PH (HO 2596/1-1), and Austrian Science Foundation (FWF-I1346) to HH). In addition, financial support by the DFG – SFB 1309 – 325871075 (to EMH and MG) and the Young Scholars’ Program of the Bavarian Academy of Sciences and Humanities (to EMH) is acknowledged. AAB thanks for financial support from the DFG-funded Cluster of Excellence Balance of the Microverse (EXC 2051). We acknowledge funding from the European Community’s Seventh Framework Program (FP7/2007-2013) under BioStruct-X (grant agreement N°283570).

## COMPETING INTERESTS

The authors declare that they have no conflict of interest.

## Supplementary Data

**Figure S1.**
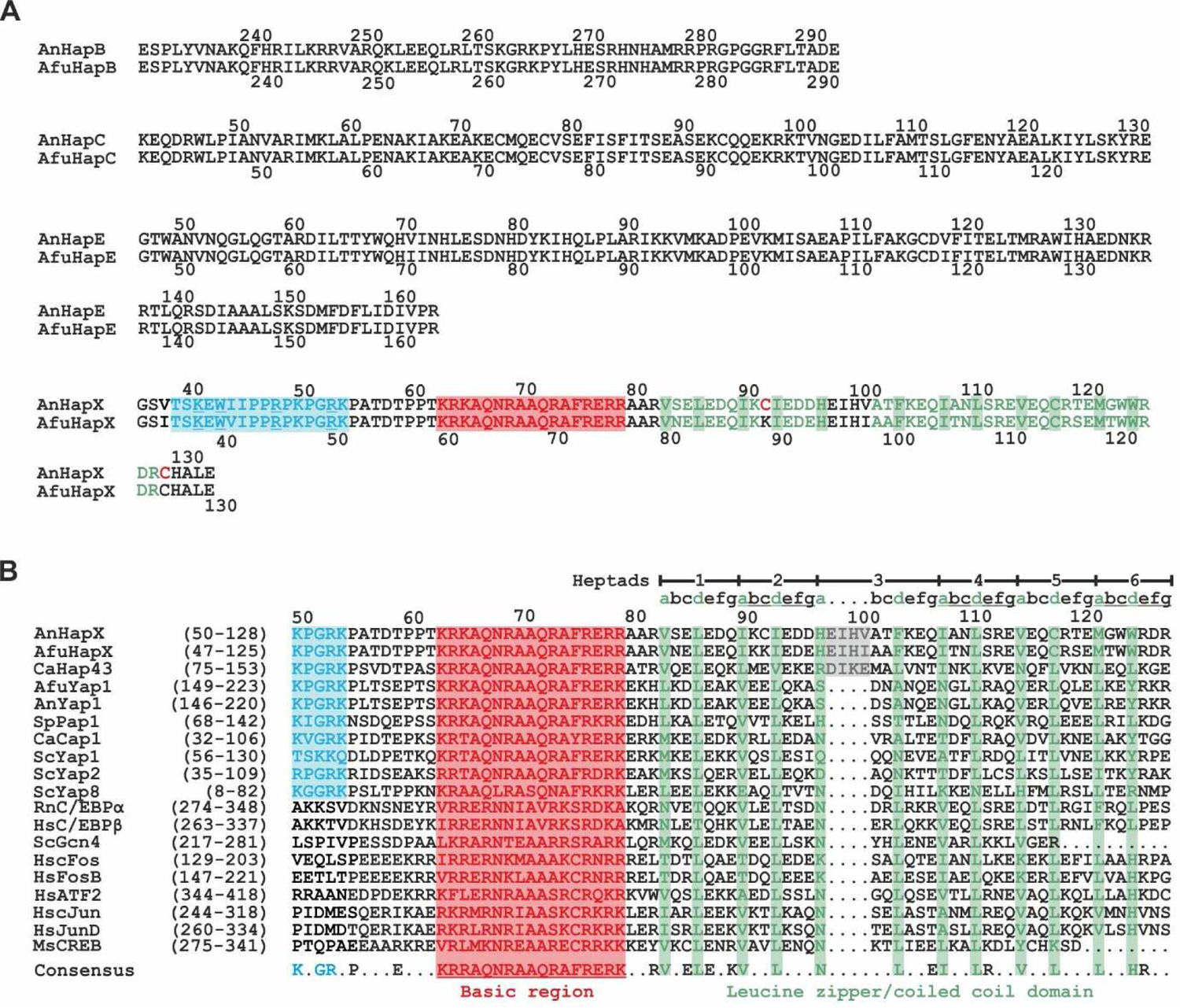
Sequence alignments. (A) Sequence alignments of *Aspergillus nidulans* (An) and *Aspergillus fumigatus* (Afu) HapB, HapC, HapE and HapX core subunits. HapX residues colored blue belong to the Hap4-like (Hap4L) domain, interacting with the CBC and DNA. The basic region of HapX, rich in lysine and arginine residues and required for DNA binding is red, while the zipper region with the characteristic hydrophobic residues of each heptad repeat is colored green. Cysteine residues of AnHapX that were mutated for crystallization are highlighted red. Residues of the Hap4L domain that were chosen for structure-based mutagenesis are underlined. (B) Sequence alignment of basic regions and coiled-coil domains of An/AfuHapX subunits as well as related bZIP factors from *Candida albicans* (Ca), *Schizosaccharomyces pombe* (Sp), *Saccharomyces cerevisiae* (Sc), *Rattus norvegicus* (Rn), *Homo sapiens* (Hs) and *Mus musculus* (Ms). Leucine zipper heptads are colored green. Note the insertion of four residues in heptad three in HapX subunits (highlighted gray) compared to other bZIP transcription factors. Color coding is according to panel (A).

**Figure S2.**
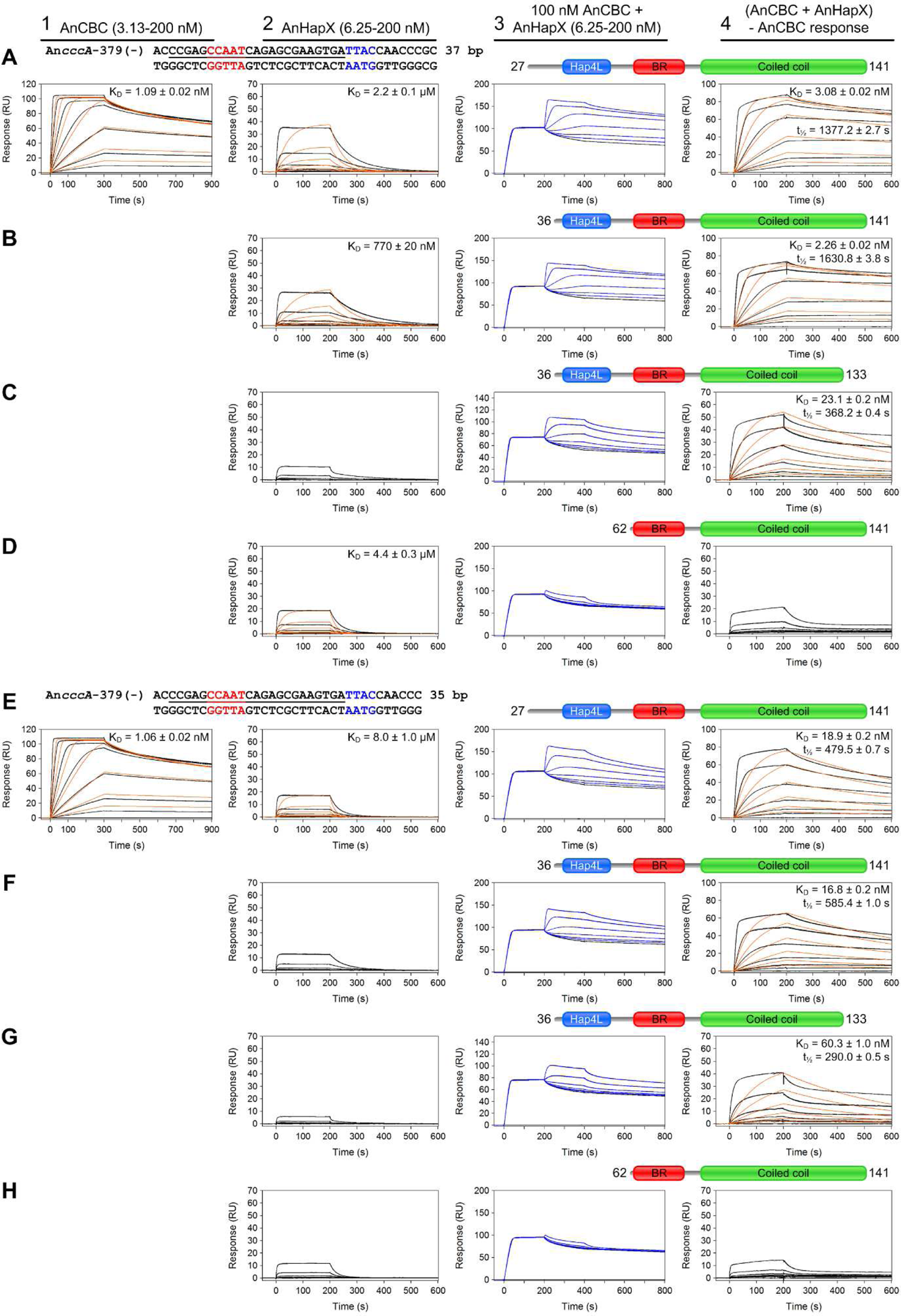
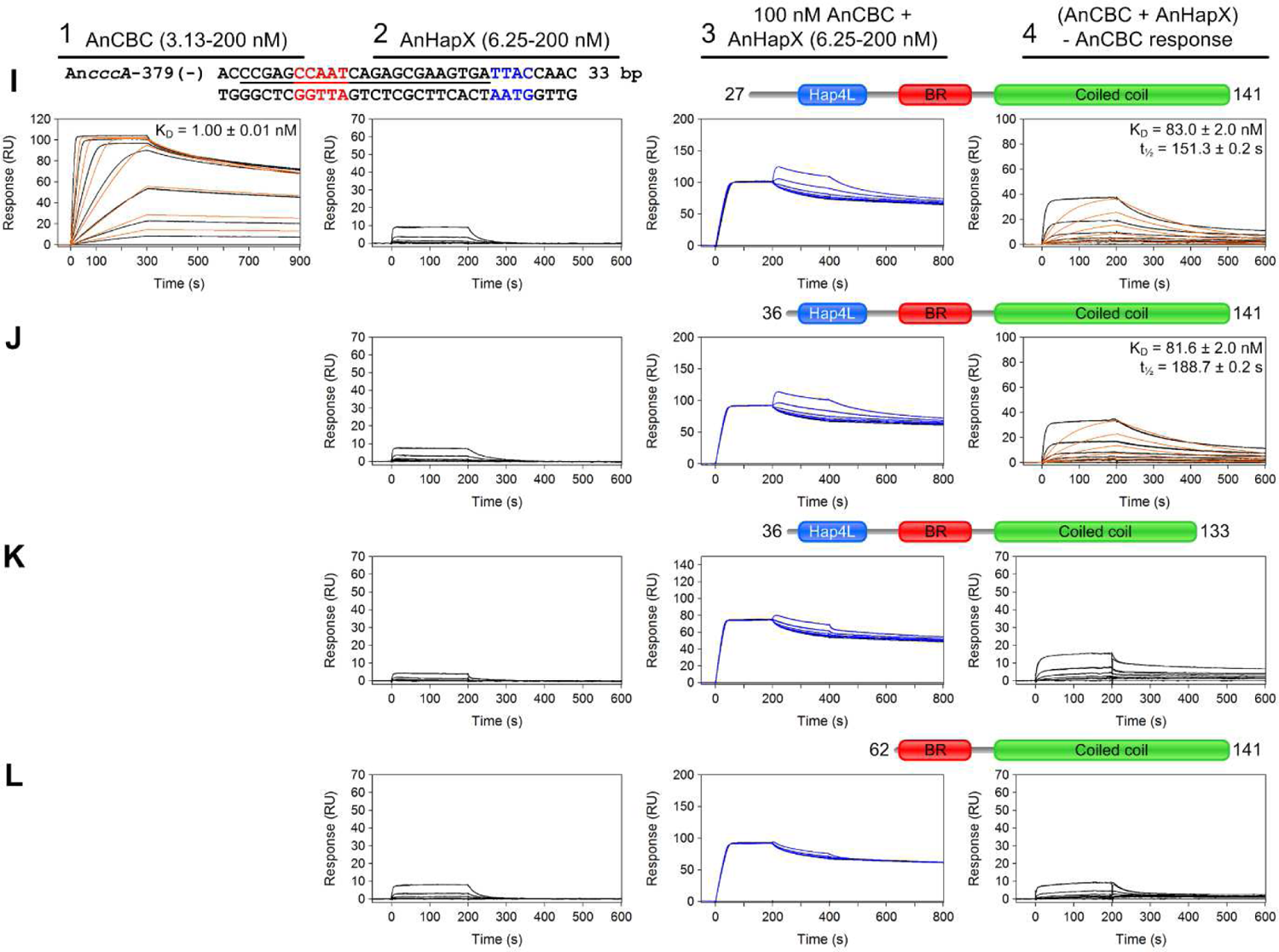
Determination of the minimal DNA duplex length and HapX protein size for cooperative CBC-HapX binding. SPR co-injection analysis of binding of different HapX constructs [(A, E, I) HapX^27-141^, (B, F, J) HapX^36-141^, (C, G, K) HapX^36-133^ and (D, H, L) HapX^62-141^ as a negative control] to preformed CBC-DNA complexes. Duplexes of various lengths [(A-D) 37 bps, (E-H) 35 bps, (I-L) 33 bps] were derived from the An*cccA* promoter. Protein sequences originate from *A. nidulans*. SPR analyses included binding of the CBC to DNA (panel 1), HapX to DNA (panel 2) and HapX to preformed CBC-DNA complexes (panel 3). Nucleotides (nt) underlined in black are covered by the CBC and nt marked in blue represent the HapX consensus motif (Furukawa et al., 2020). Binding responses of the indicated CBC or HapX concentrations injected in duplicate (black lines) are overlaid with the best global fit derived from a 1:1 interaction model including a mass transport term (red lines). Binding responses of CBC-DNA-HapX ternary complex formation (panel 3, blue lines) were obtained by concentration dependent co-injection of HapX on preformed binary CBC-DNA complexes after 200 seconds within the steady-state phase. Sensorgrams in panel 4 depict the association/dissociation responses of HapX on preformed CBC-DNA, and were obtained by CBC response subtraction (co-injection of buffer) from HapX co-injection responses. Dissociation constants (K_D_) and half-lives of the complexes including their standard deviation are plotted inside the graphs.

**Figure S3.**
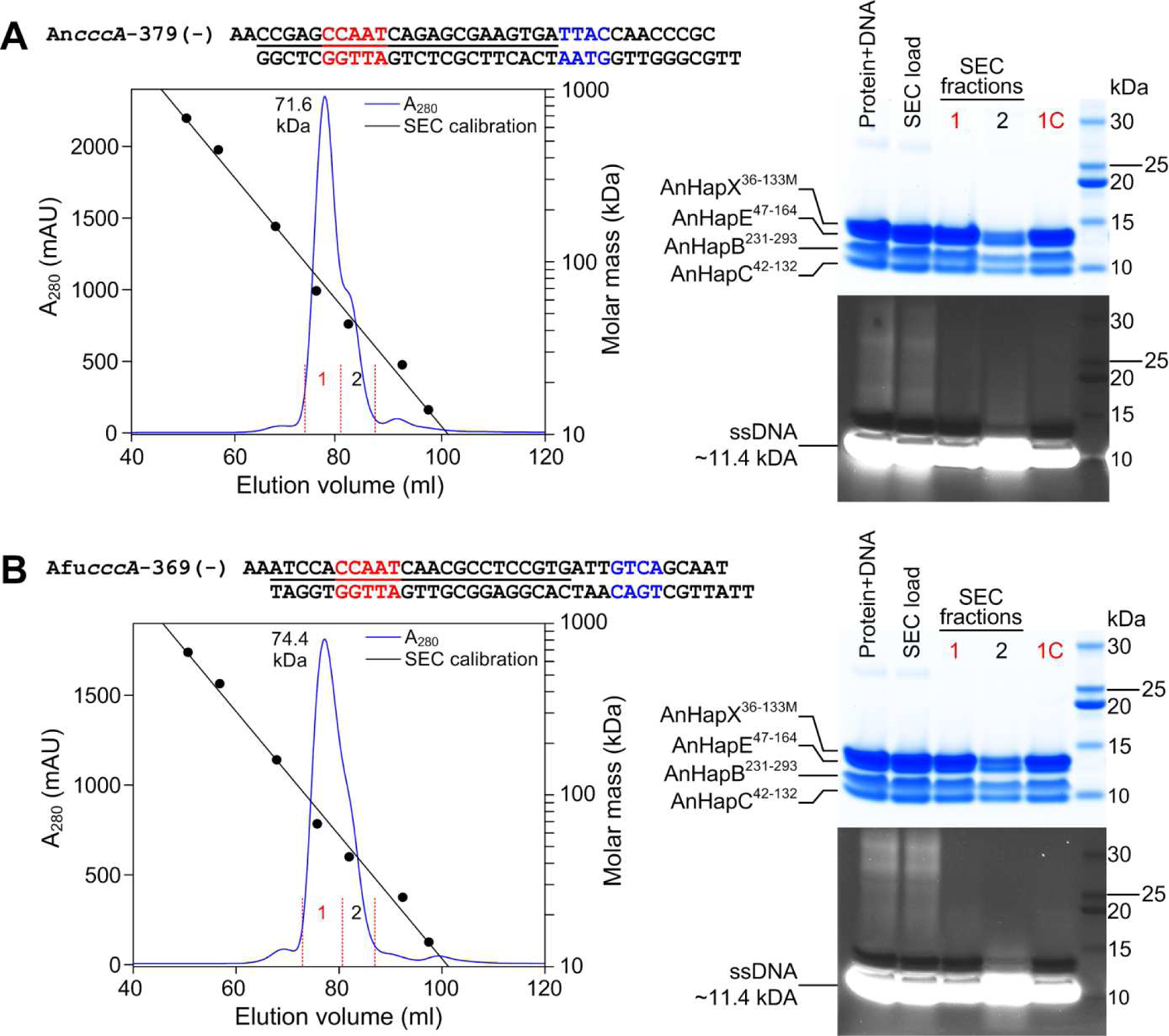
Reconstitution of CBC-DNA-HapX complexes for crystallization. (A, B) Size exclusion chromatography (SEC) profiles and SDS-PAGE analysis of prepared CBC-HapX-DNA complexes from *A. nidulans*. The high affinity CBC-HapX target sites from both *A. nidulans* (A) and *A. fumgiatus cccA* promoter sequences were chosen as DNA duplexes. SEC fractions that were subjected to crystallization after a subsequent concentration step are marked in red (samples labeled 1C in the respective SDS-PAGE gels). SDS-PAGE gels were stained for protein with the GelCode Blue Stain Reagent (upper panels) prior to DNA staining with SYBR Gold Nucleic Acid Gel Stain (lower panels). Source data are provided as separate files.

**Figure S4.**
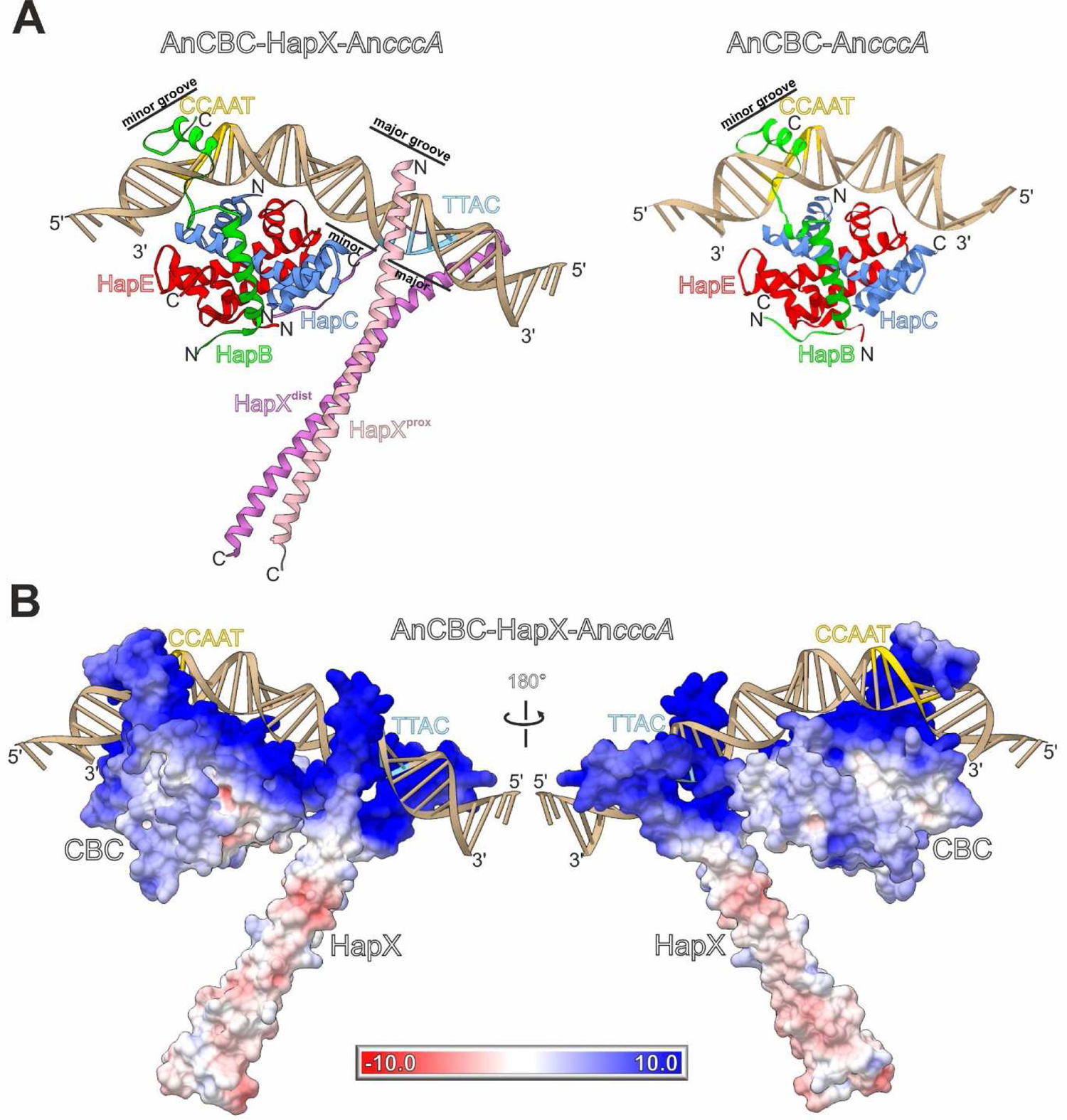
CBC-HapX complex bound to double stranded DNA. (A) Ribbon illustration of the *A. nidulans* CBC (PDB ID 6Y37 (Hortschansky et al., 2020)) and CBC-HapX structures bound to DNA derived from the *A. nidulans cccA* promoter. Binding of HapX does not induce any structural rearrangements within the core CBC. Color-coding is according to Figure 2. (B) Connolly surface representations of the AnCBC-HapX-An*cccA* complex with colors indicating positive (blue, 10 kT/e) and negative (red, −10 kT/e) electrostatic surface potentials.

**Figure S5.**
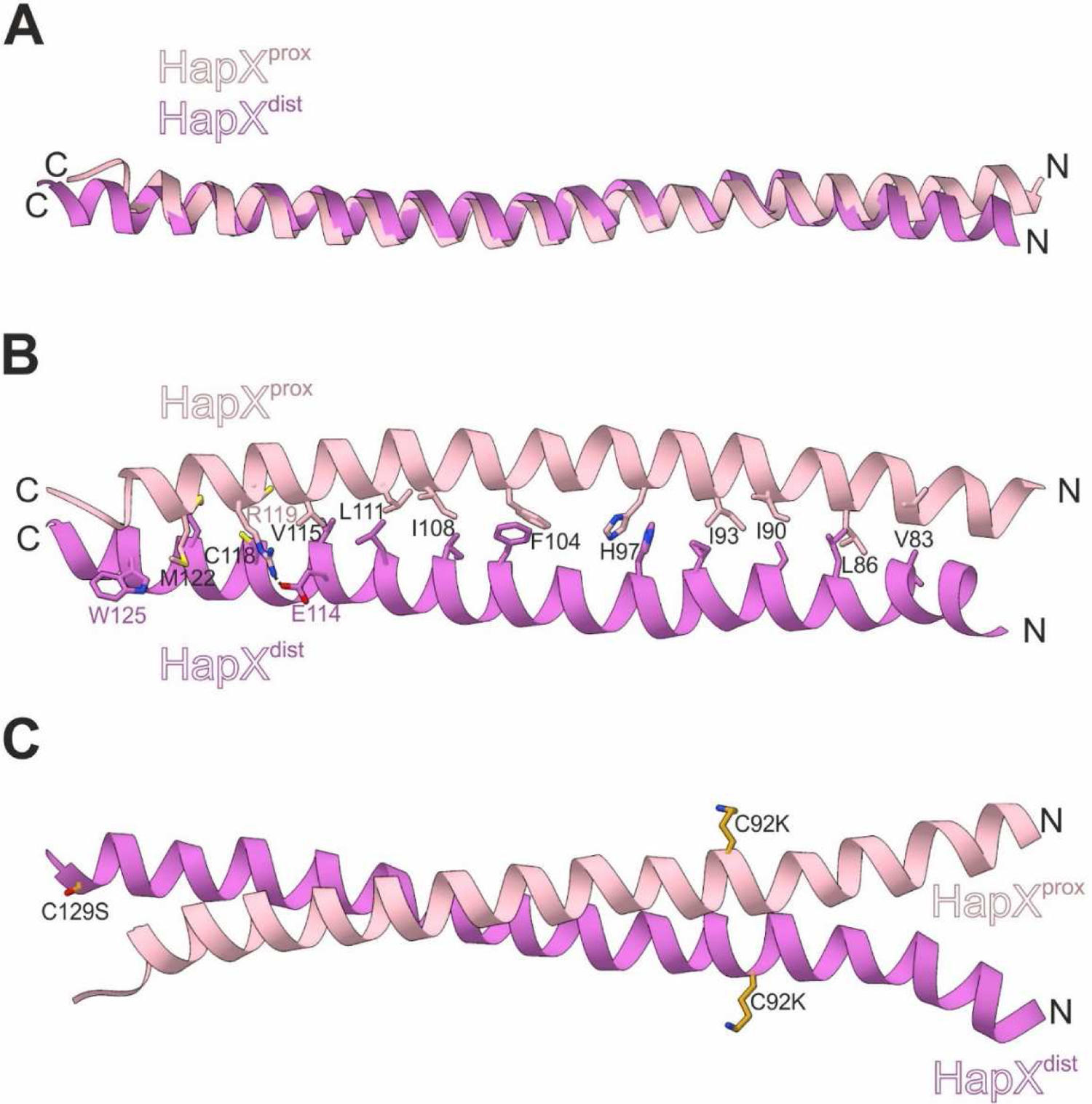
HapX subunit structure and interactions. (A) Superposition of full helical segments of HapX subunits illustrates their structural similarity. (B) Amino acids involved in hydrophobic and polar contacts within the coiled-coil segment (residues 77-130) are shown as sticks and labelled by the one-letter code. Residues identical in both chains are marked black, while individual ones are labelled magenta or light pink. (C) Mutations C92K and C129S of HapX have been engineered to prevent disassembly of the CBC-HapX complex due to oxidation of the cysteine residues. The sites of mutations are surface-exposed and not engaged in protein-protein interactions.

**Figure S6.**
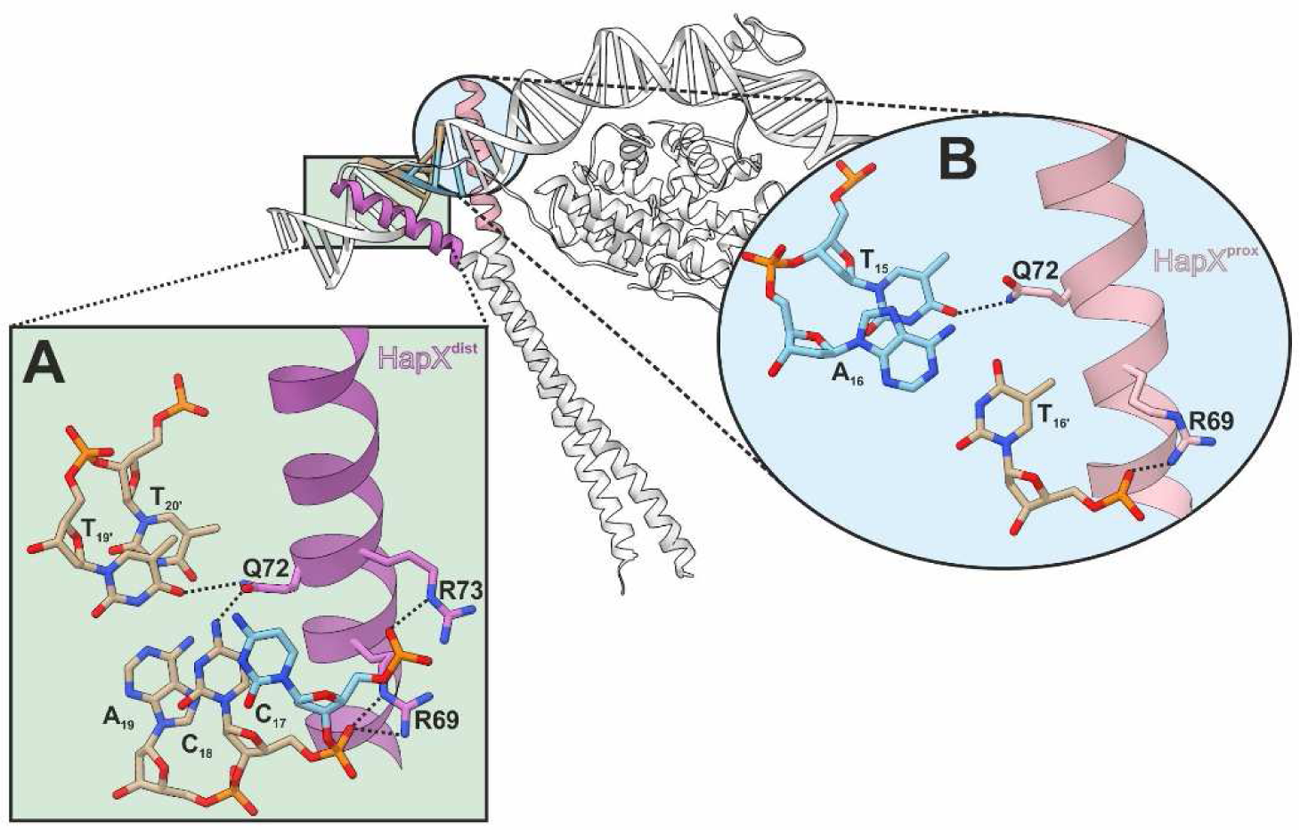
Interaction of the bZIP domains of HapX with DNA major grooves. Illustration of key polar interactions of HapX^dist^ (A) and HapX^prox^ (B) with the major groove. The interactions are mapped on the overall structure of the protein-DNA complex in the middle (see also Figures S7E, F and S8).

**Figure S7.**
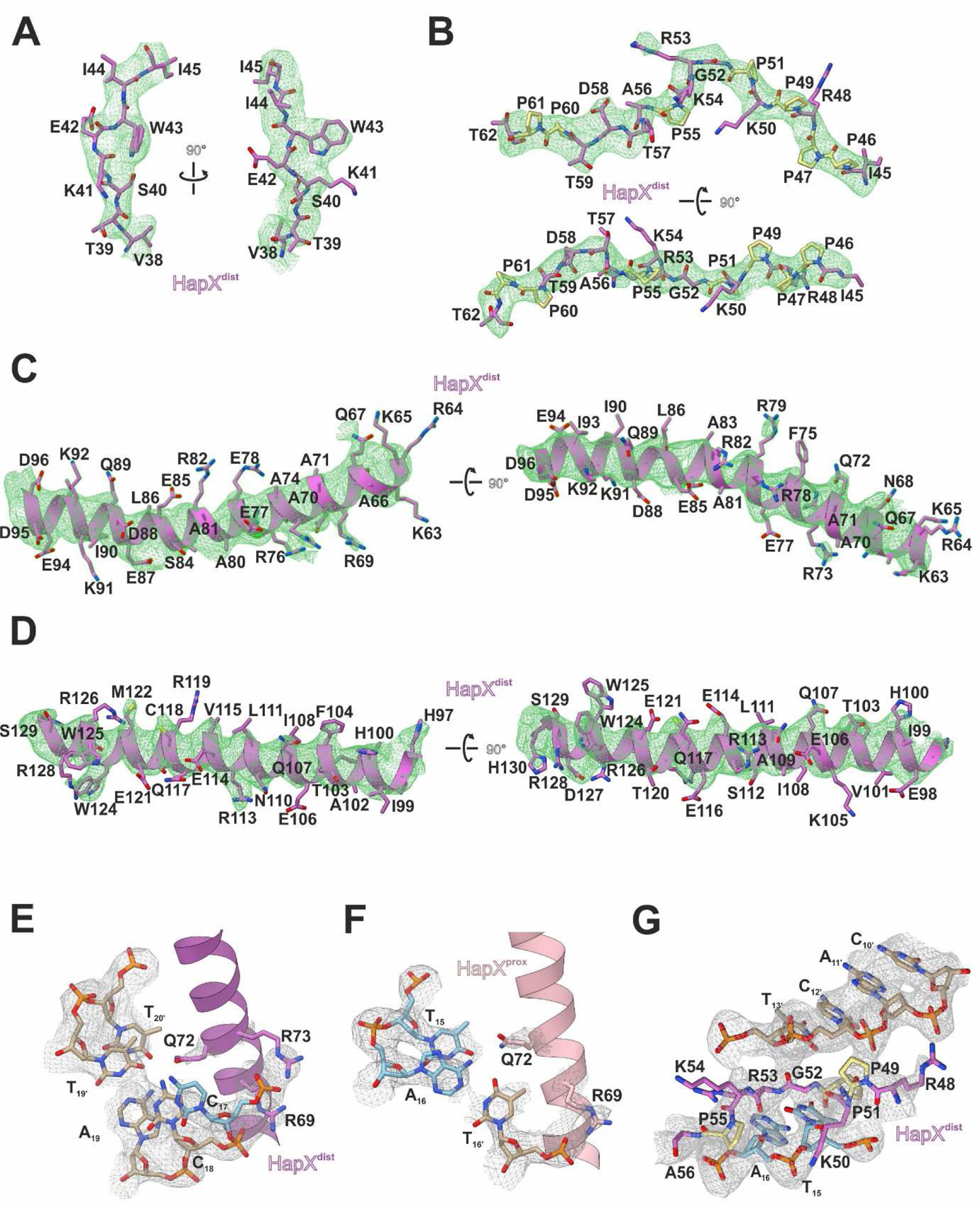
Electron density maps for the CBC-HapX-An*cccA* complex. (A-D) F_O_-F_C_ omit electron density maps (green meshes contoured to +3 σ) for various sections of HapX^dist^ confirm the presence of full-length HapX^dist^ in the crystal lattice. The displayed structural parts were omitted from the model prior to phasing. The well-defined residues Trp43 (A) Arg69 (C) and Trp124 (D) were crucial for sequence assignment. (E-G) 2F_O_-F_C_ electron density maps (gray meshes contoured to 1 σ) are shown for the interaction of HapX^dist^ (E, see Figure S6A) and HapX^prox^ (F, see Figure S6B) with the DNA major groove as well as for HapX^dist^ with the minor groove (G, see Figure 2D).

**Figure S8.**
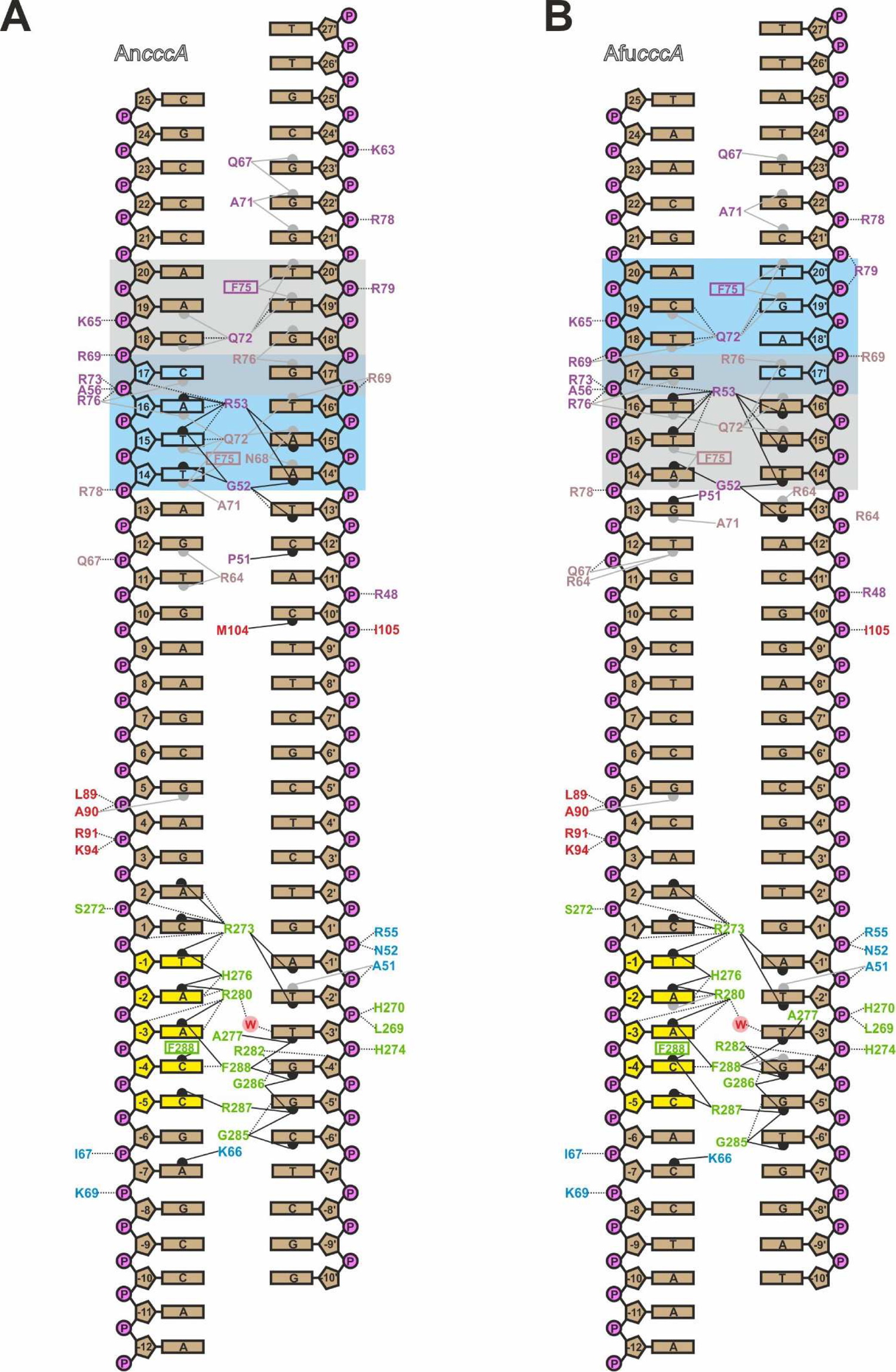
Schematic illustration of protein-DNA contacts. Interactions of the *A. nidulans* CBC-HapX complex with *cccA* DNA from (A) *A. nidulans* and (B) *A. fumigatus*. Hydrogen bonds are indicated by black dotted lines. Amino acids stacked with bases are boxed. Color coding is according to Figure 2. Minor and major groove contacts, determined with DNAproDB (Sagendorf et al., 2017; Sagendorf et al., 2020), are shown by black respectively gray half circles and connection lines. The consensus and nonconsensus half-sites of the pseudopalindromic HapX interaction motif are shaded in blue and gray, respectively. Interactions of the core AnCBC with *cccA* were taken from the high-resolution structure PDB ID 6Y37 (Hortschansky et al., 2020).

**Figure S9.**
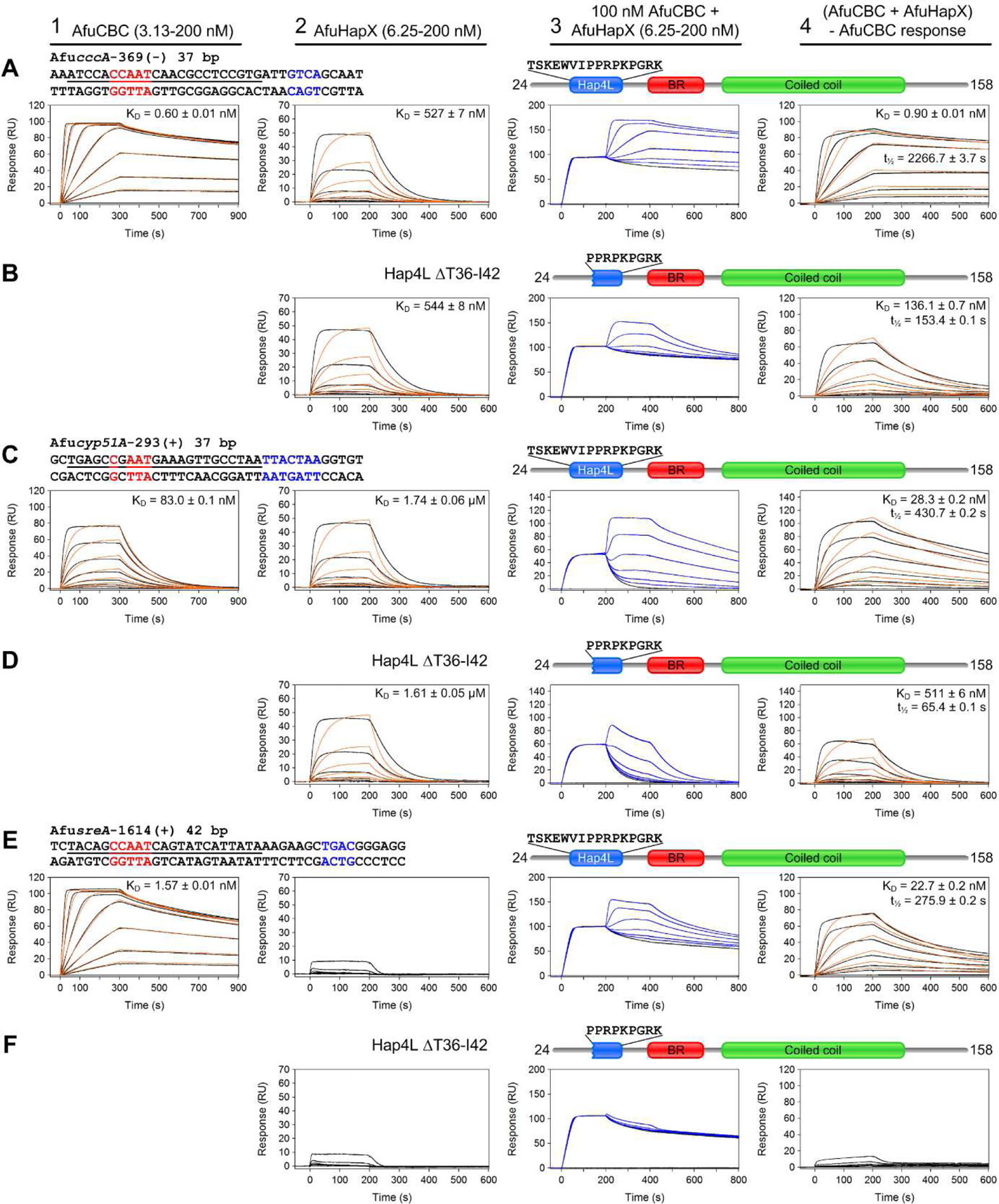
Effects of partial deletion (residues 36-42) of the Hap4L domain of AfuHapX on DNA binding. SPR co-injection analysis of binding of wt and mutant AfuHapX^24-158^ ΔT36-I42 proteins to preformed AfuCBC-DNA complexes [(A, C, E) wt HapX, (B, D, F) HapX ΔT36-I42]. DNA duplexes were derived from Afu*cccA* (A-B), Afu*cyp51A* (C-D) and Afu*sreA* (E-F) promoter sequences. Proteins originate from *A. fumigatus*. Data are presented as described in the legend to Figure S2.

**Figure S10.**
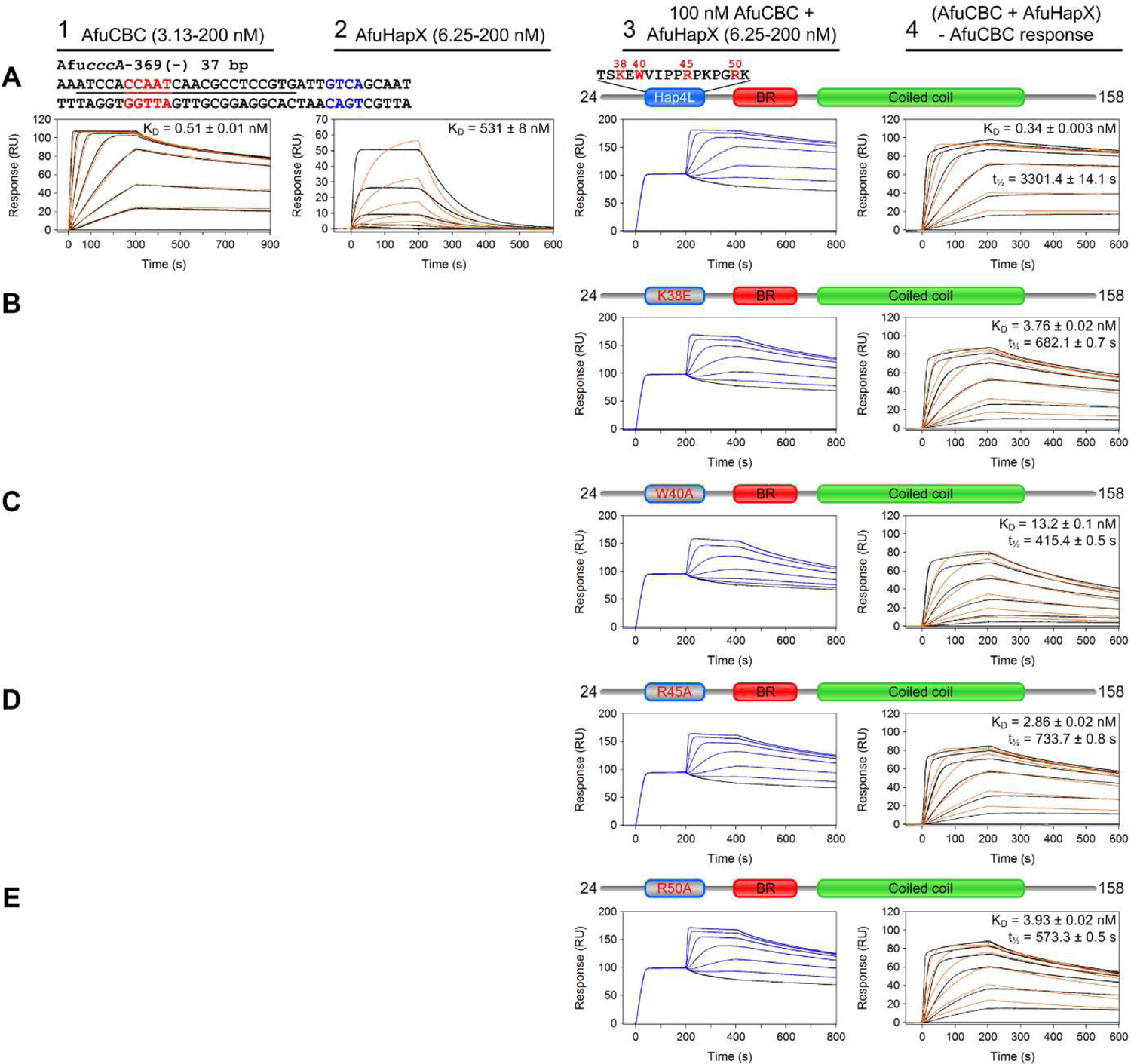
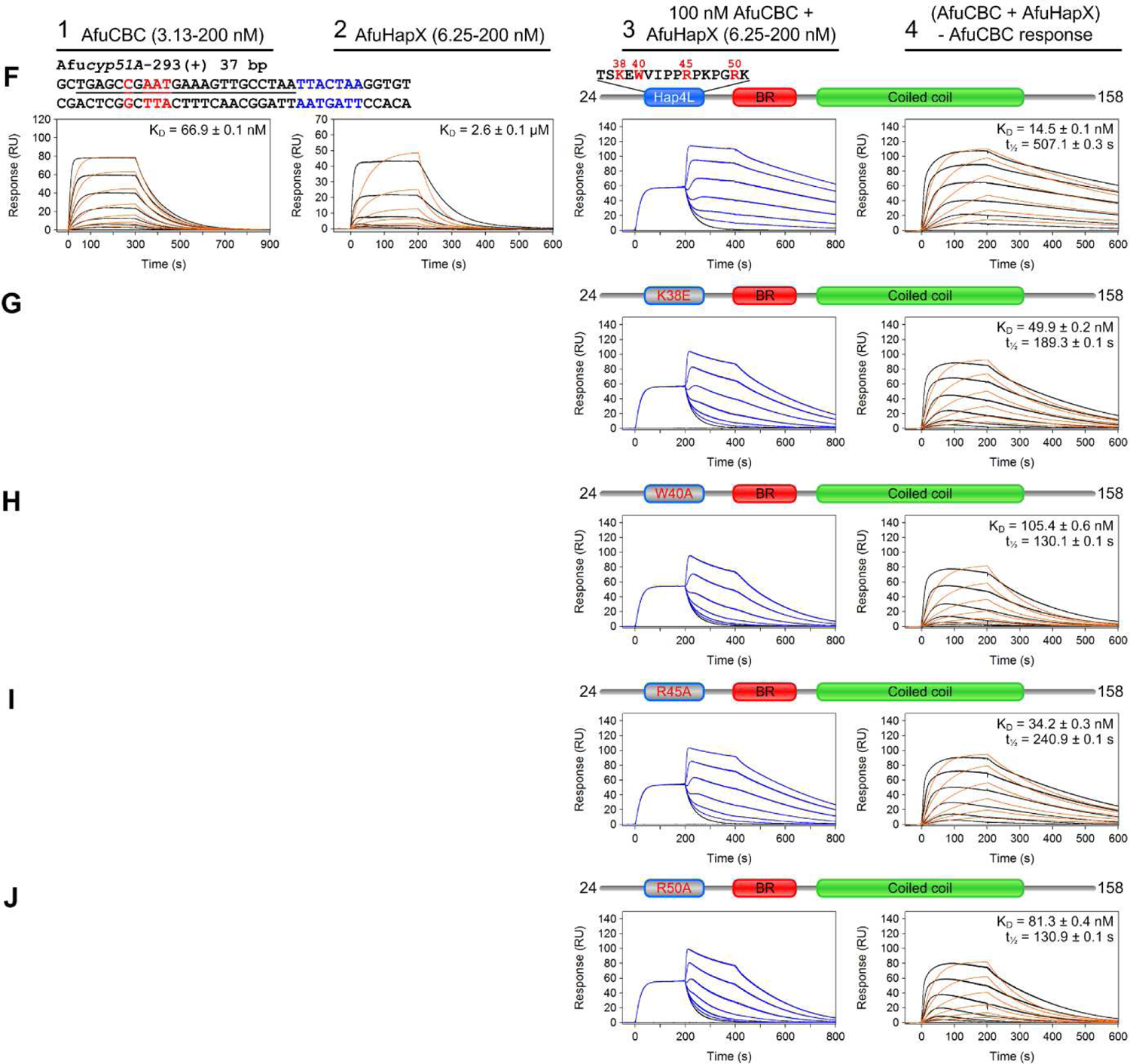
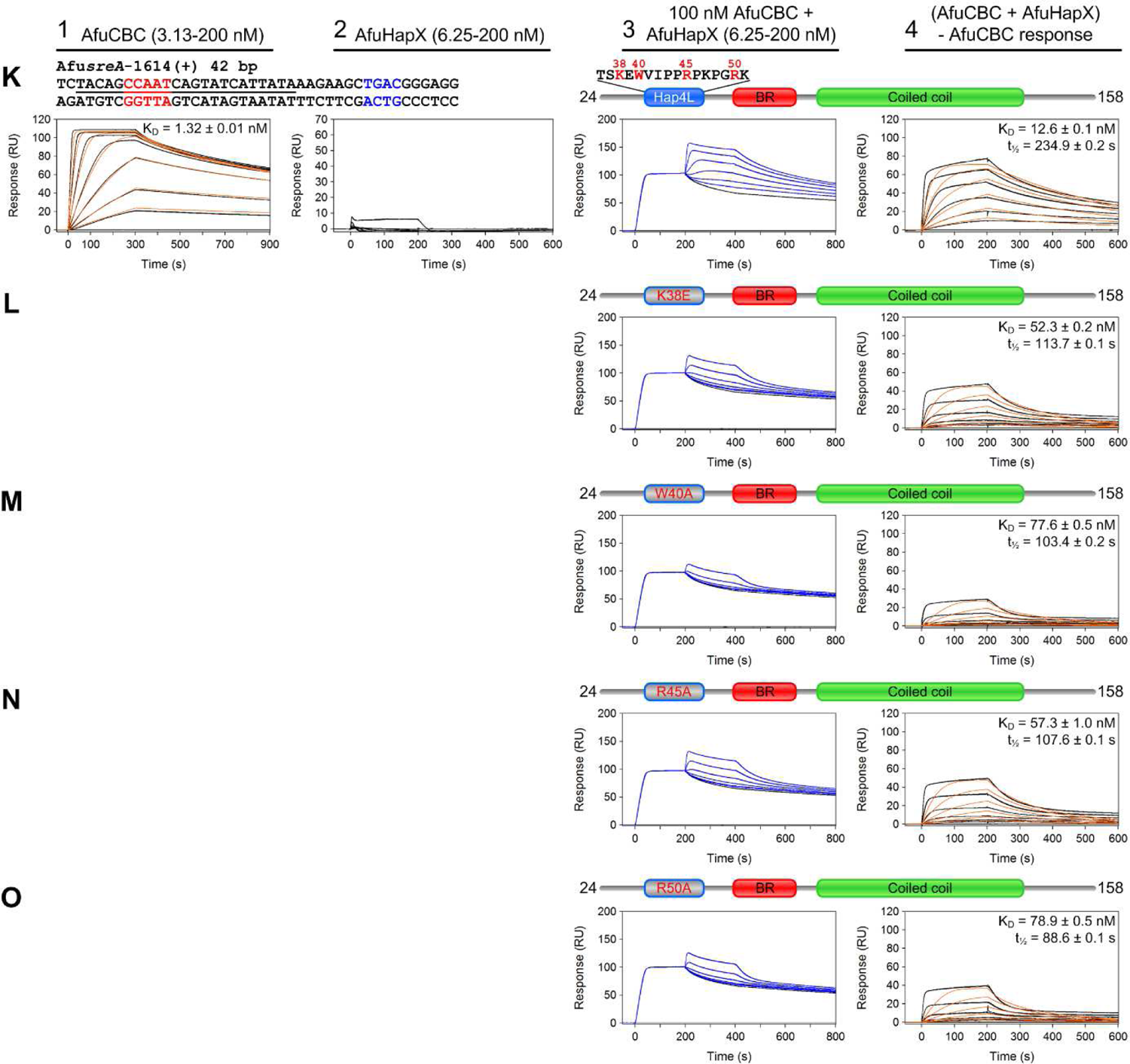
Effects of AfuHapX Hap4L domain mutations on DNA binding. SPR co-injection analysis of binding of wt and mutant AfuHapX^24-158^ proteins to preformed AfuCBC-DNA complexes [(A, F, K) wt HapX, (B, G, L) HapX^K38E^, (C, H, M) HapX^W40A^, (D, I, N) HapX^R45A^ and (E, J, O) HapX^R50A^]. DNA duplexes were derived from Afu*cccA* (A-E), Afu*cyp51A* (F-J) and Afu*sreA* (K-O) promoter sequences. Proteins originate from *A. fumigatus*. Data are presented as described in the legend to Figure S2.

**Figure S11.**
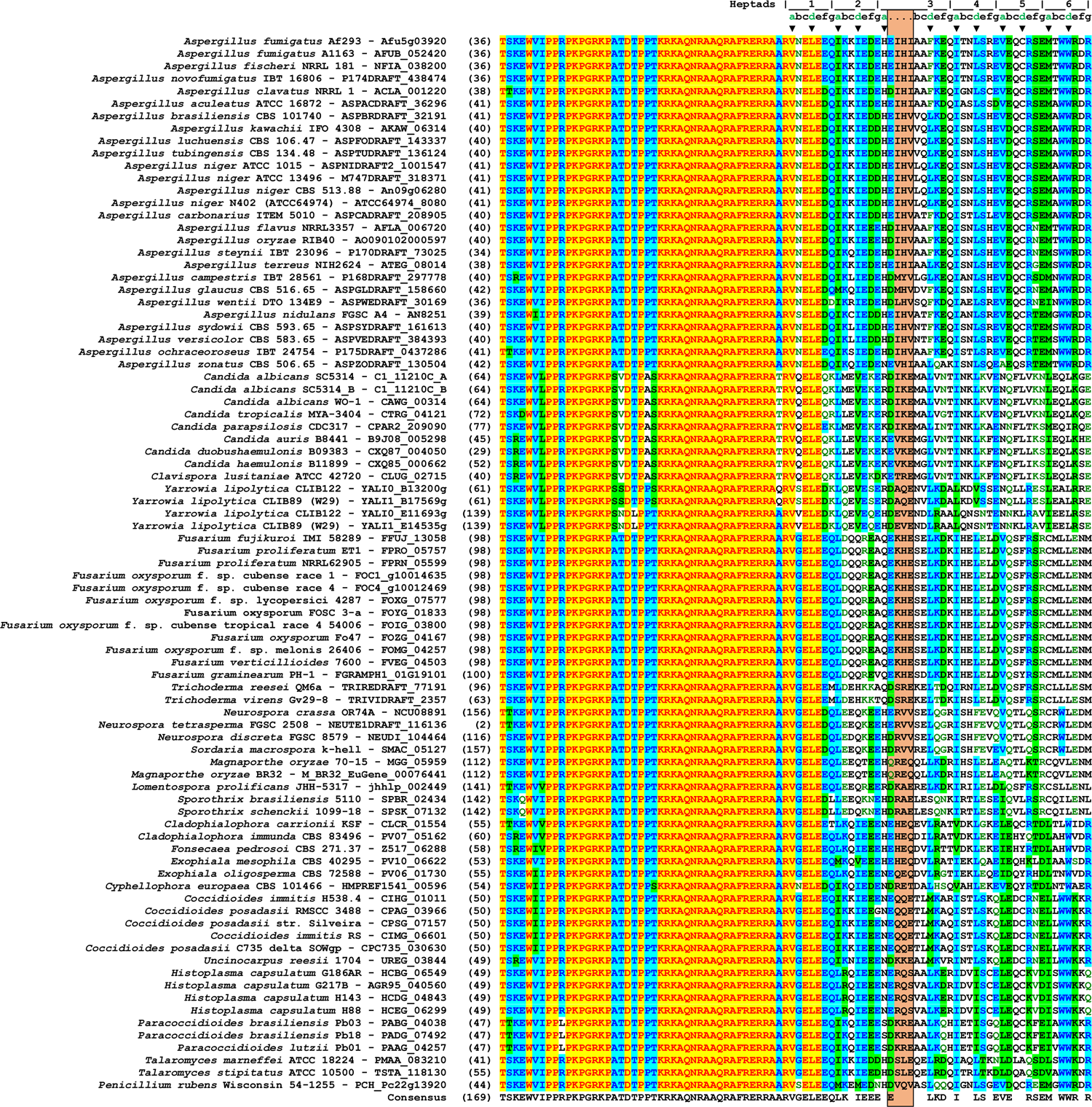
Sequence alignment of the Hap4L domain and bZIP region of HapX subunits from various organisms. Each sequence is preceded by its source organism and its gene ID. Identical residues are marked in yellow, residues conserved in 50% of the sequences are shaded in light blue and blocks of similar residues are marked in green. The insertion of four residues in heptad three is highlighted in orange. Alignments were performed with AlignX (Vector NTI Advance 11). Sequences included represent all HapX subunits that were deposited in the publicly accessible database FungiDB (https://fungidb.org (Basenko et al., 2018)) on the 5^th^ of February 2021.

**Figure S12.**
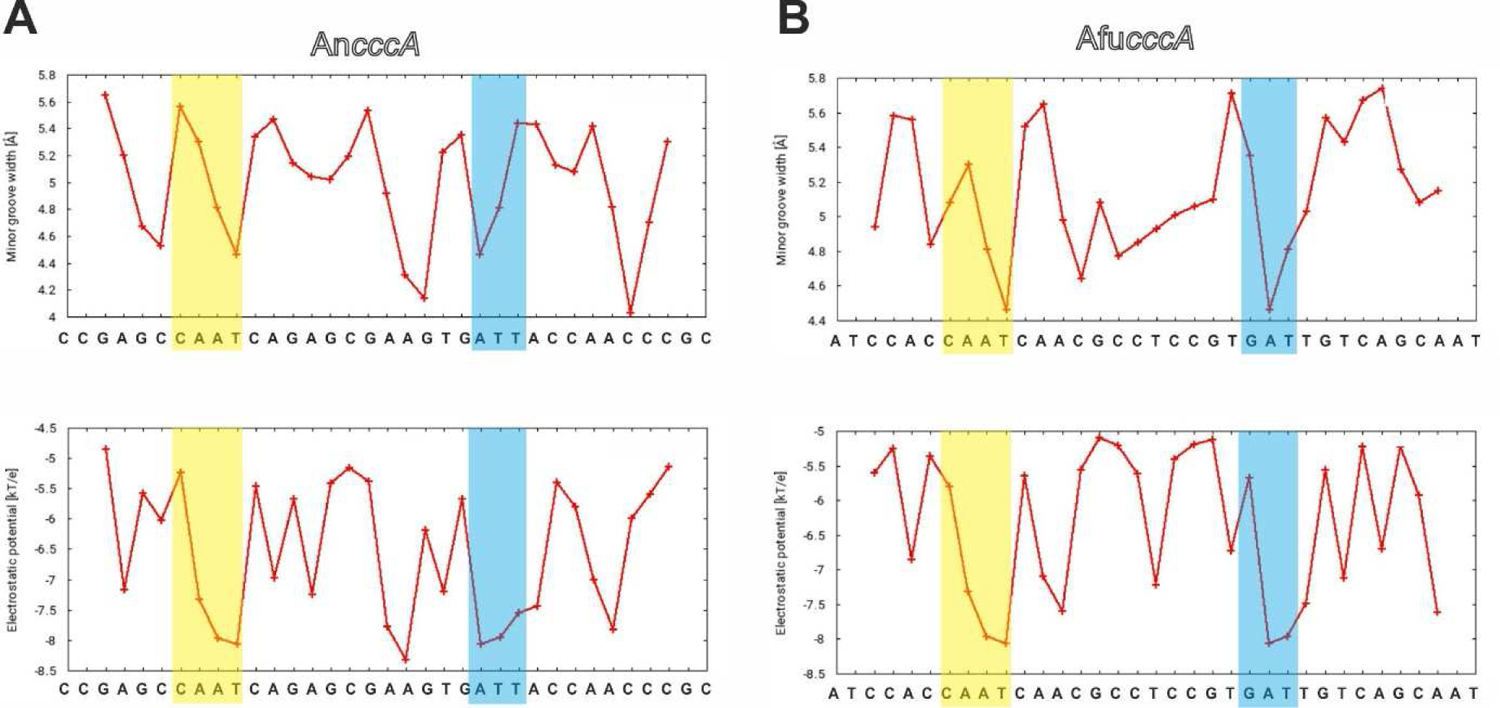
Analysis of DNA shape and electrostatic potential. (A, B) *A. nidulans* (A) and *A. fumigatus* (B) *cccA* promoter sequences were analyzed with the webtool DNAphi for minor groove width and electrostatic potential. Narrow minor grooves (< 5 Å width (Rohs et al., 2009)) are associated with more negative electrostatic potential and have a preference to bind arginine residues. For the CBC-HapX-DNA complex, Arg53 of HapX^dist^ targets the narrow minor groove at the AT-rich region (blue shaded) downstream of the CCAAT box (yellow).

**Figure S13.**
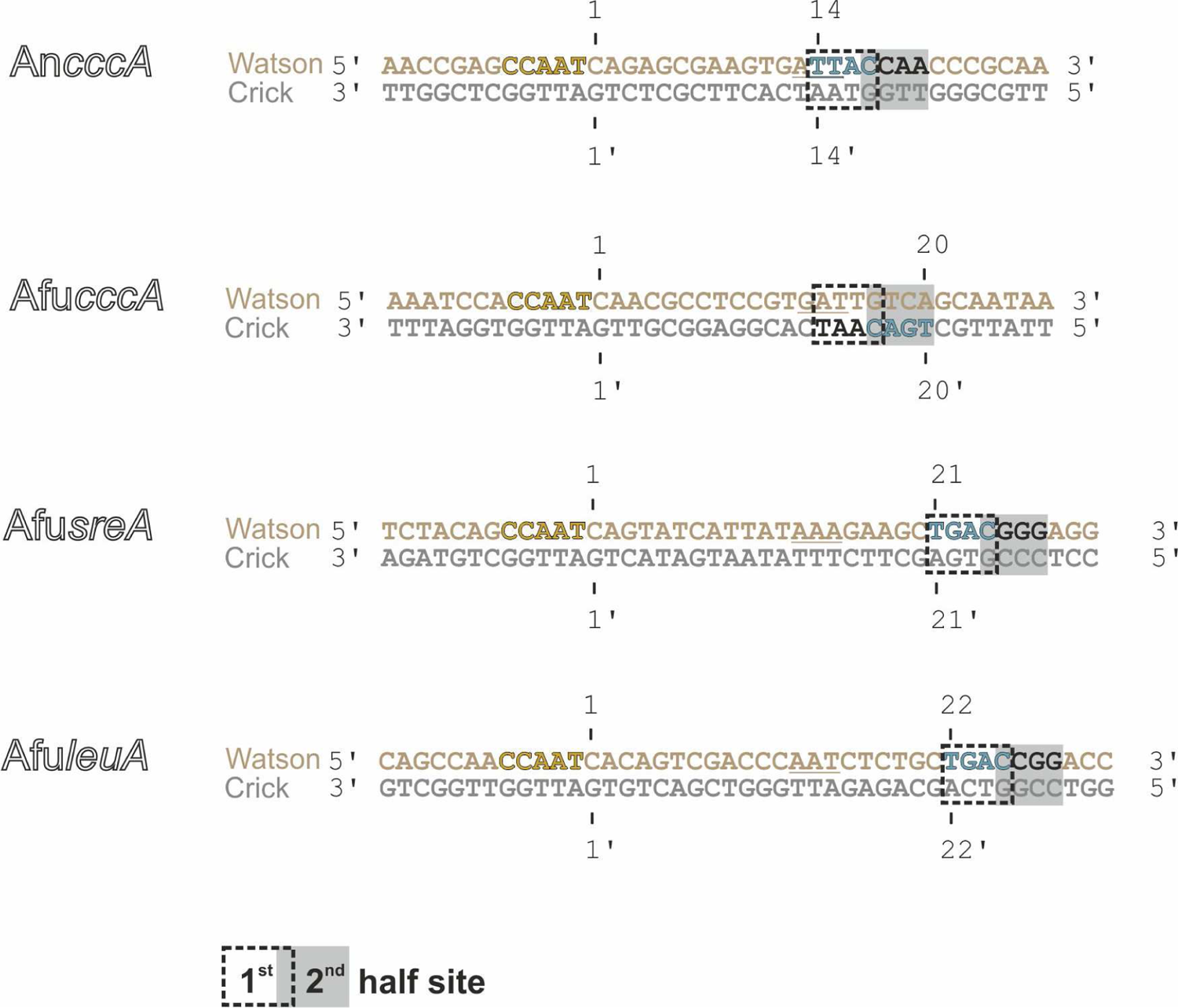
Sequence comparison of selected promoter DNA fragments used for crystallization and SPR analysis. The HapX recognition site TTAC/TGAC is located either on the Watson (position 14; An*cccA*) or on the Crick (position 20; Afu*cccA*) strand. Promoter sequences of *sreA* and *leuA* genes show an unusually large distance between CCAAT and HapX binding sites that are both located on the Watson strand. In fact their spacing is 7 respectively 8 bases larger than for An*cccA*. Underlined nucleotides represent the AT-rich region (RWT submotif) targeted by the Hap4L domain of HapX.

**Figure S14.**
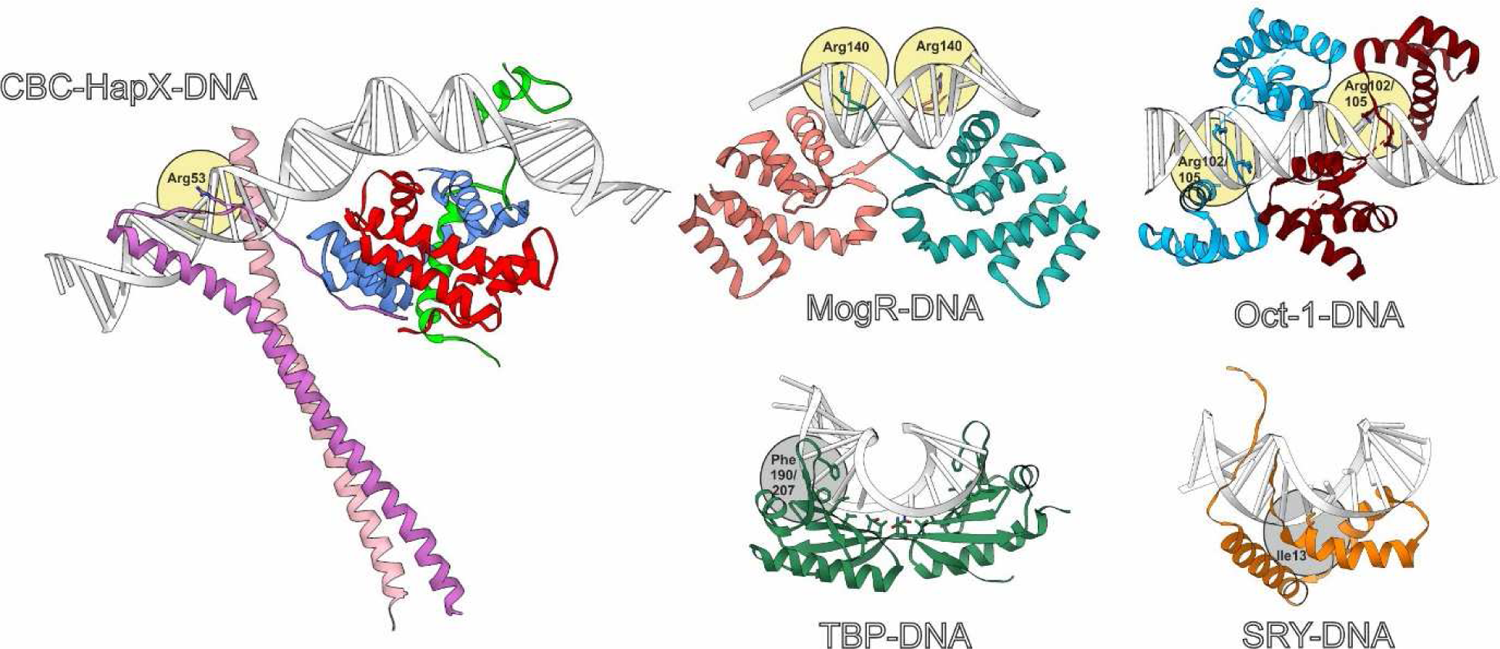
Comparison of transcription factors targeting the minor groove. Structures of the CBC-HapX complex (PDB ID 7AW7), the MogR repressor (PDB ID 3FDQ (Shen et al., 2009)), the Oct-1 transcription factor (PDB ID 1HF0 (Remenyi et al., 2001)), the TATA-box binding protein (TBP; PDB ID 1YTB (Kim et al., 1993)) and the sex-determining region Y protein SRY (PDB ID 1J46 (Murphy et al., 2001)) bound to DNA. HapX, MogR and Oct-1 recognize DNA minor groove shape by arginine residues (highlighted by yellow circles), whereas TBP and SRY transcription factors associate with minor grooves *via* hydrophobic contacts (key amino acid residues are highlighted by gray circles and labelled by the three-letter-code).

**Table S1.** X-ray data collection and refinement statistics.

|  | AnCBC-HapX-AncccA | AnCBC-HapX-AfucccA |
| --- | --- | --- |
| **Crystal parameters** |  |  |
| Space group | I222 | I222 |
| Cell constants | a= 76.1 Å<br>b= 133.1 Å<br>c= 199.8 Å | a= 76.8 Å<br>b= 127.7 Å<br>c= 199.0 Å |
| Subunits / AU<sup>a</sup> | 1 CBC<br>2 HapX<br>1 DNA duplex | 1 CBC<br>2 HapX<br>1 DNA duplex |
| **Data collection** |  |  |
| Beam line | X06SA, SLS | X06SA, SLS |
| Wavelength (Å) | 1.0 | 1.0 |
| Resolution range (Å)<sup>b</sup> | 47-3.4<br>(3.5-3.4) | 47-3.5<br>(3.6-3.5) |
| No. observations | 74959 | 55853 |
| No. unique reflections<sup>c</sup> | 14326 | 12478 |
| Completeness (%)<sup>b</sup> | 99.6 (99.9) | 97.9 (99.1) |
| R<sub>merge</sub> (%)<sup>b, d</sup> | 5.8 (52.7) | 3.8 (66.4) |
| I/σ (I)<sup>b</sup> | 14.7 (2.8) | 18.5 (2.1) |
| **Refinement (REFMAC5)** |  |  |
| Resolution range (Å) | 30-3.4 | 30-3.5 |
| No. refl. working set | 13590 | 11837 |
| No. refl. test set | 716 | 623 |
| No. non hydrogen | 5053 | 5033 |
| Solvent (Na<sup>+</sup>, Cl<sup>-</sup>) | 3 | 1 |
| R<sub>work</sub>/R<sub>free</sub> (%)<sup>e</sup> | 25.2/29.7 | 24.5/29.1 |
| r.m.s.d. bond/angle (Å) / (°)<sup>f</sup> | 0.002/1.022 | 0.002/1.151 |
| Average B-factor (Å<sup>2</sup>) | 136.6 | 173.0 |
| Ramachandran Plot (%)<sup>g</sup> | 96.9/2.9/0.2 | 98.3/1.7/0.0 |
| PDB accession code | 7AW7 | 7AW9 |

[a] Asymmetric unit
[b] The values in parentheses for resolution range, completeness, R<sub>merge</sub> and I/σ (I) correspond to the highest resolution shell
[c] Data reduction was carried out with XDS and from a single crystal. Friedel pairs were treated as identical reflections
[d] $R_{\text{merge}}(I) = \sum_{hkl} \sum_j |I(hkl)_j - \langle I(hkl) \rangle| / \sum_{hkl} \sum_j I(hkl)_j$ , where $I(hkl)_j$ is the $j^{\text{th}}$ measurement of the intensity of reflection hkl and $\langle I(hkl) \rangle$ is the average intensity
[e] $R = \sum_{hkl} | |F_{\text{obs}}| - |F_{\text{calc}}| | / \sum_{hkl} |F_{\text{obs}}|$ , where R<sub>free</sub> is calculated without a sigma cut off for a randomly chosen 5% of reflections, which were not used for structure refinement, and R<sub>work</sub> is calculated for the remaining reflections
[f] Deviations from ideal bond lengths/angles
[g] Percentage of residues in favored region / allowed region / outlier region

**Table S2.** A. fumigatus strains used in this study

| Strain | Background | Genotype | Reference |
| --- | --- | --- | --- |
| AfS77 (wt) | ATCC 46645 | $\Delta akuA::loxP$ | (Hartmann et al., 2010) |
| $\Delta hapX$ | AfS77 | $\Delta hapX::ptrA$ | (Gsaller et al., 2014) |
| *venus* *hapX* | AfS77 | $\Delta hapX::ptrA$, *venus* *hapX*, *hph* | (Gsaller et al., 2014) |
| *venus* *hapX*<sup>K38E</sup> | AfS77 | $\Delta hapX::ptrA$, *venus* *hapX*<sup>K38E</sup>, *hph* | this study |
| *venus* *hapX*<sup>W40A</sup> | AfS77 | $\Delta hapX::ptrA$, *venus* *hapX*<sup>W40A</sup>, *hph* | this study |
| *venus* *hapX*<sup>R50A</sup> | AfS77 | $\Delta hapX::ptrA$, *venus* *hapX*<sup>R50A</sup>, *hph* | this study |

**Table S3.**
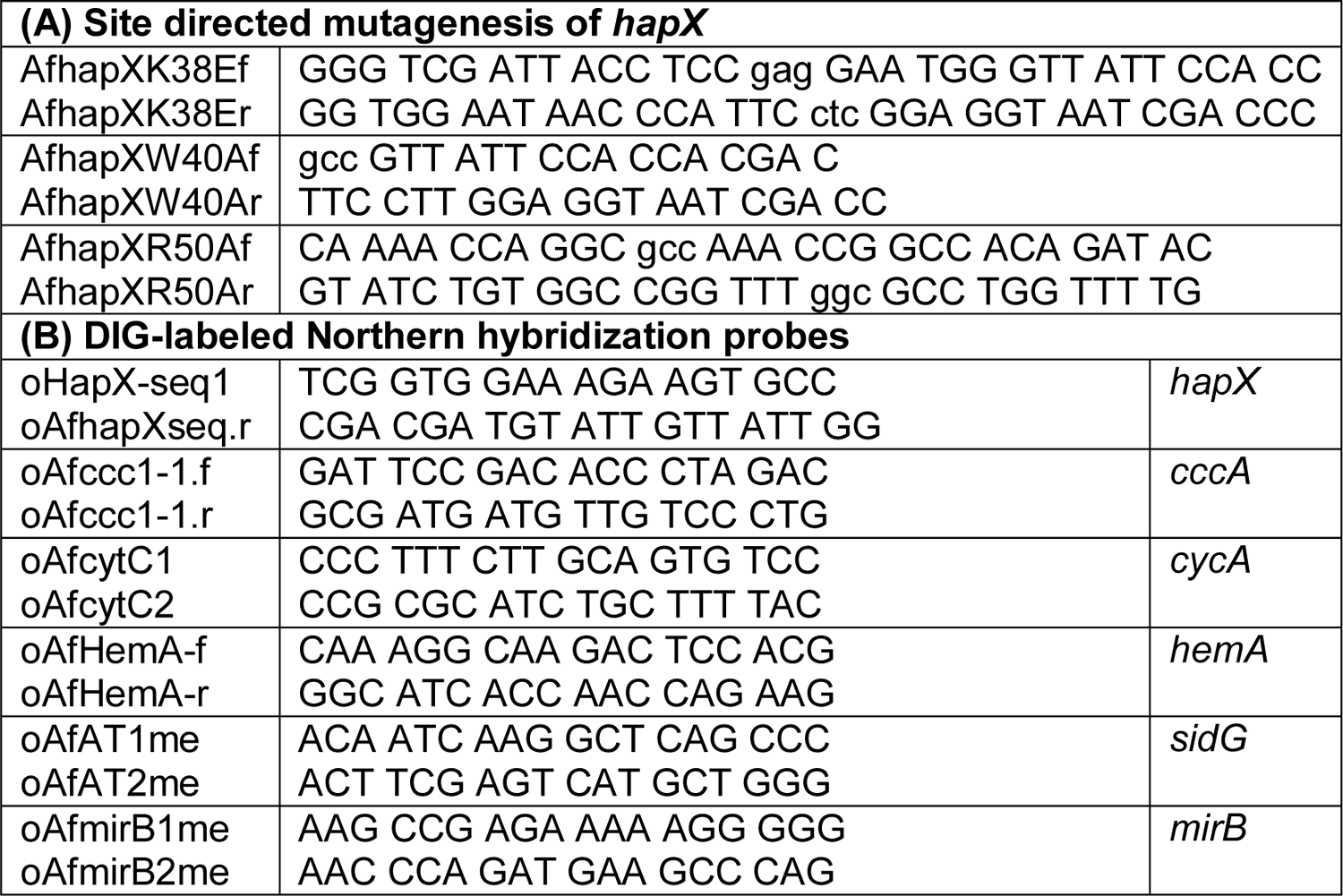
Oligonucleotides used in this study (5’→3’)

## Notes

### Competing Interest Statement

The authors have declared no competing interest.

